# Storing >1 byte of information in 16S ribosomal RNA using orthogonal trans-splicing ribozymes

**DOI:** 10.64898/2026.07.14.738544

**Authors:** Matthew J. Dysart, Lin Fang, Lavanya Karinje, James Chappell, Lauren B. Stadler, Jonathan J. Silberg

## Abstract

Catalytic-RNA (cat-RNA) expressed from mobile DNA can record cellular events, such as the uptake of plasmids via horizontal gene transfer, by splicing a barcode onto 16S ribosomal RNA (rRNA) - a system termed RNA addressable modification (RAM). However, scaling RAM to record multiple simultaneous biological events requires large numbers of orthogonal cat-RNA whose signals reflect the biological features under investigation rather than variability arising from the barcode sequence. Here, we explore how to design orthogonal cat-RNA to record information about multiple plasmid-encoded traits in parallel. We show that cat-RNA having tRNA-derived barcodes with sequence variation in the anticodon stem-loop present greater signal consistency within *Escherichia coli* than mRNA-derived barcodes. When orthogonal cat-RNA designs harboring tRNA-derived barcodes were evaluated in *Vibrio natriegens* and *Pseudomonas putida*, increased variance was observed compared with *Escherichia coli*. Nevertheless, the signal consistency was sufficient to use these orthogonal cat-RNAs to report on the relative activities of four promoters and two origins of replication by sequencing barcoded-rRNA derived from the three organisms. These results show how RAM can be multiplexed to report on mobile DNA features in microbial communities and illustrate the importance of accounting for variability in RNA outputs when designing and interpreting multiplexed RNA barcoding data.

**GRAPHICAL ABSTRACT:** 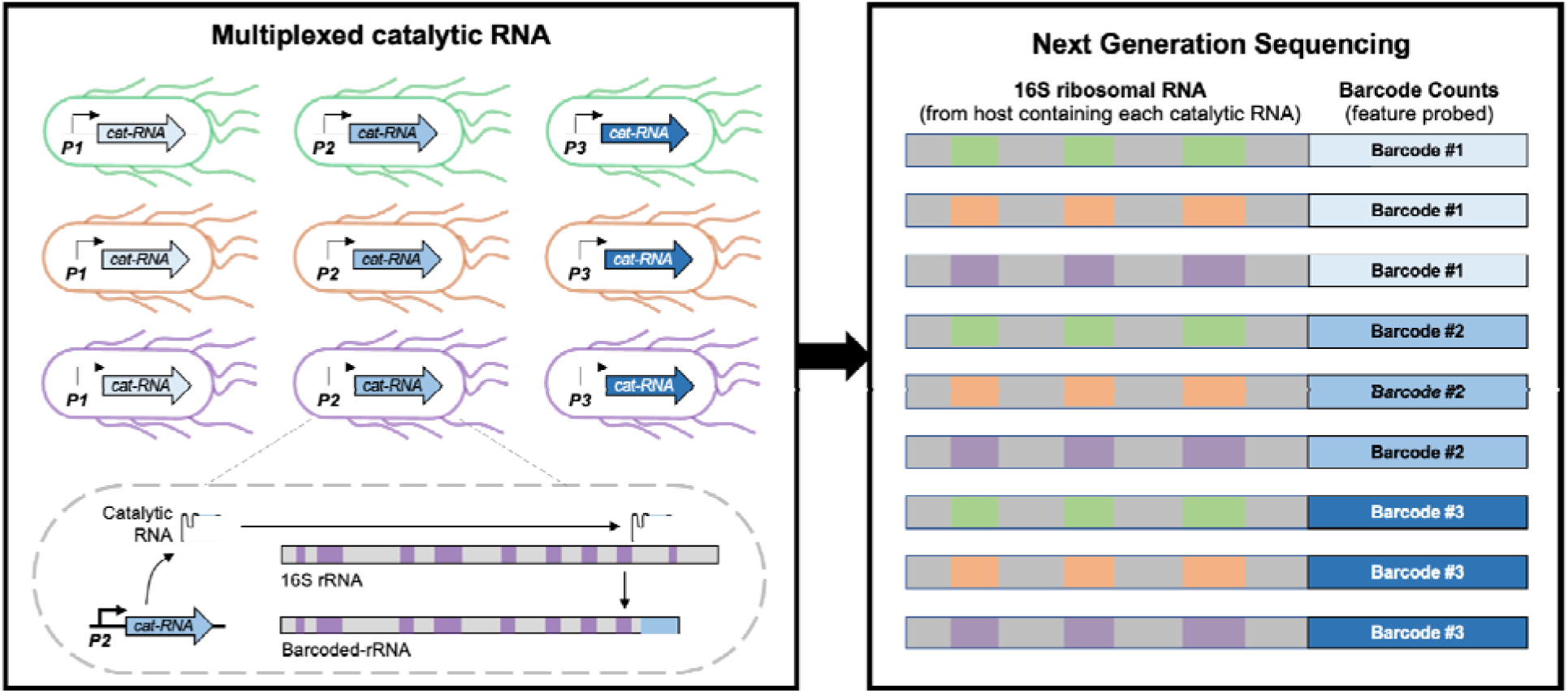

## INTRODUCTION

Genetic barcoding is widely used to link genotypes and phenotypes, enabling multiple bits of cellular information to be read out in parallel using sequencing (1–3). When leveraged for CRISPR-Cas9 screens, barcodes can be used as identifiers in DNA constructs expressing guide RNAs that target different genomic sites (4, 5). To date, DNA barcoding has been used to multiplex targeting of different loci in parallel, including CRISPR interference (6), CRISPR activation (7), and transposon sequencing (8). DNA barcoding has also been used to store information about sequential cellular events (9), for lineage tracing (10), to map genetic design space (11), and to report on gene transfer (12). RNA barcoding has also increased the throughput of cellular assays by allowing for sequencing readouts (13, 14). Massively parallel studies of transcription can now be achieved by amending barcodes to all mRNA within single cells and varying barcode indexes across each cell in a population (15, 16). Furthermore, RNA barcoding has been used to understand how changes in mRNA sequence affect stability (17).

Recently, an autonomous RNA barcoding tool was reported that can be genetically coded in cells and used to store information about which microbes in a community take up mobile DNA via conjugation and transduction (18, 19). With this approach, which is called RNA addressable modification (RAM), a catalytic RNA (cat-RNA) splices a barcode onto 16S rRNA *in vivo* (20). By sequencing barcoded-rRNA in communities, the host range of mobile genetic elements (MGEs) can be monitored across microbiomes (18, 19). RAM can also be used to monitor the host range of two MGEs in parallel (18), illustrating its potential for multiplexed measurements. However, scaling RAM to larger numbers of simultaneous measurements requires a larger set of orthogonal cat-RNAs whose signals reflect the biological features under investigation within communities rather than variability arising from differences in cat-RNA activity or barcode stability.

Nucleic acid barcodes present a range of challenges for multiplexing (21). RNA sequence modifications can differentially affect transcript stability (22, 23), translation initiation (24, 25), and catalytic activity (26, 27). Also, mRNA stability is influenced by the sequence and length of the 5′ untranslated region, RNAase-binding motifs, interactions with translation machinery, and small molecules (28, 29). While the controls on RNA synthesis and degradation have been intensively studied to understand their regulation (30), the impact of small sequence modifications on the steady-state levels of synthetic RNA remains poorly defined in bacteria. Recently, rational engineering of the barcode sequence of RAM, specifically incorporating a structured tRNA scaffold rather than an mRNA fragment, was shown to increase and stabilize the barcoded-rRNA signal (31).

To better understand how to apply RAM for multiplexed information storage in bacteria, we investigated how varying the cat-RNA barcode affects the steady-state levels of barcoded-rRNA produced in *Escherichia coli*. We benchmark the signal generated by cat-RNA that amend orthogonal barcodes derived from a mRNA fragment (Figure 1), designated cat-RNA-v1, which was used in the prototype RAM system (18), and show that these present greater variation than cat-RNA having orthogonal barcodes built using a structured tRNA, designated cat-RNA-v2, which was recently reported (31). We also evaluate the performance of cat-RNA-v2 across *E. coli*, *Vibrio natriegens*, and *Pseudomonas putida* and show that it can be multiplexed to report on four promoters and two origins of replication.

**Figure 1.**
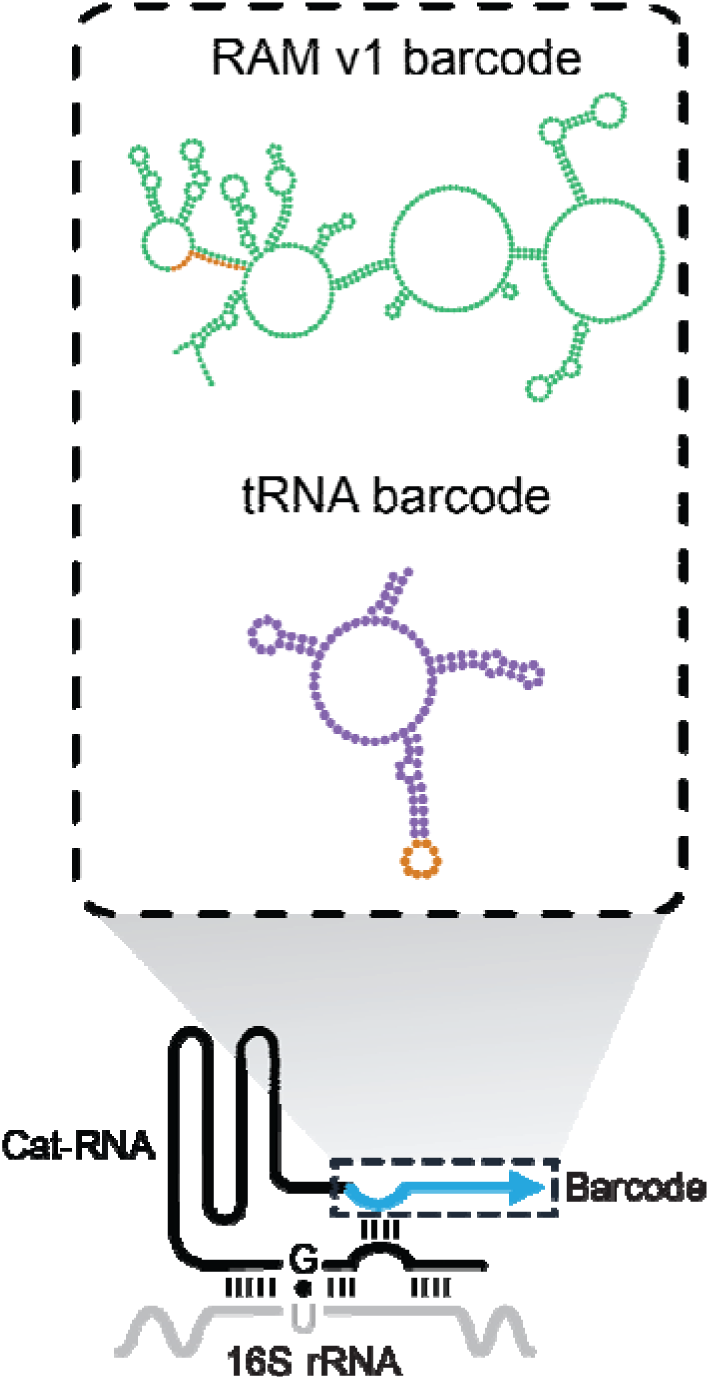
Using RAM to report on biological features in cells. (*bottom*) The cat-RNA contains a ribozyme with a guide that targets splicing to 16S rRNA and a barcode (blue). *(top*) For cat-RNA-v1, the barcode is composed of a gfp fragment (green) and a thirteen base pair insert (orange) representing the orthogonal internal barcode. Cat-RNA-v2 uses a tRNA-derived barcode (purple). With cat-RNA-v2, a ten base pair nucleotide substitution in the anti-Shine-Dalgarno middle stem-loop (orange) is varied to create orthogonal barcodes.

## MATERIALS AND METHODS

### Reagents

Primers were synthesized by Millipore Sigma (Burlington, Massachusetts). Enzymes for cloning from New England Biolabs (Ipswich, MA) included Q5® High-Fidelity DNA Polymerase (M0491S), T4 DNA Ligase (M0202S), T4 polynucleotide kinase (M0201S), BsaI-HFv2 (R3733S), and BbsI-HF (R3539S). Phanta Max Super-Fidelity DNA Polymerase as part of the 2 × Phanta Max Master Mix Dye Plus (P525-01) was from Vazyme (San Diego, California), while Esp3I (ER0451) was from Thermo Fisher Scientific (Waltham, MA). All enzymes used recommended buffers. Reagents for RNA extractions were from Thermo Fisher Scientific and included the MagMAX™ Microbiome Ultra Nucleic Acid Isolation Kit (A42357), Microbiome Lysis Solution (A42361), Viral/Pathogen Proteinase K (A42363), Viral/Pathogen Elution Buffer (A42364), TURBO DNAse (AM2239), and DNAse buffer (AM1907). RT-qPCR reagents were from PCR Biosystems (Wayne, Pennsylvania) and Thermo Fisher Scientific, which included the dye-based qPCRbio Sygreen 1-step kit (PB25.11), the 20x RTase Go reverse transcriptase with RNAse inhibitors kit (PB10.53-05), and TaqMan™ Fast Virus 1-Step Master Mix (4444434). Kits for DNA extractions were from Qiagen (Germantown, Maryland) and Promega (Madison, WI) and included the QIAprep Spin Miniprep Kit (27104) and Wizard® Genomic DNA Purification Kit (A1120). RT-PCR reactions were from New England Biolab, including LunaScript® Multiplex One-Step RT-PCR Kit (E3010), NEB Q5® High-Fidelity DNA Polymerase (M0491S), NEB Q5®Reaction Buffer Pack (B9027), and New England Biolab’s Monarch® Spin DNA Gel Extraction Kit (T1120S). Kanamycin (30772) was from Research Products International Corp. (Mt. Prospect, IL), LB Broth (11-120) was from Apex Bioresearch Products (El Cajon, CA), and magnesium sulfate heptahydrate (M2773-500G) and potassium phosphate dibasic (P3786-500G) were from Sigma-Aldrich (St. Louis, MO). Agar (BP1423-500), magnesium chloride (P33-500), potassium chloride (P217-500), sodium chloride (S9625-500G), and glucose (A16828.36) were all from Thermo Fisher Scientific, as well as the KingFisher Apex System used for RNA extractions and the QuantStudio 5 Real-Time PCR System used for RT-qPCR.

### Biological Resources

*E. coli* MegaX DH10B T1^R^ Electrocomp™ Cells (C640003) were from Thermo Fisher Scientific (Waltham, MA), *Vibrio natriegens* Vmax (ATCC14048) and *Pseudomonas putida* F1 (ATCC700007) were from was from ATCC (Manassas, VA). Supplemental Table 1 describes all plasmids. All cat-RNA-v1 plasmids were derived from pPK076 (18), which amends a fragment of the *gfp* gene to 16S rRNA, while all cat-RNA-v2 plasmids were derived from pLNK007 (31), which transcribes a cat-RNA with the restored P1-P10 loop of the wildtype ribozyme and a barcode that is derived from a consensus *Escherichia coli* tRNA sequence.

### Statistical Analyses

To evaluate the variance in sequencing data, the Euclidean distance was calculated from each barcode element to every other barcode element (Supplemental Figure 1). Since there is only a single plane examined, the Euclidean distance simplifies to the absolute value of the difference between two given elements. Examined as a set of Euclidean distances, the magnitude between the resulting 1^st^ and 4^th^ quartiles represents the dynamic range introduced by the set of barcodes within the given organism. Finally, the sum of squares between the Euclidean distance dataset is taken, calculated through the anova1() Matlab function and reported as the MSE term. This value is used to compare the noise of a given orthogonal barcode set between conditions and/or species. Further, this value can be used to predict the number of degenerate barcodes required to observe biological differences with the given set of orthogonal barcodes by considering the limit of detection (LOD) as:

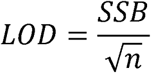

where *SSB* is the sum of the squares between (SSB) barcode distances and *n* is the number of orthogonal barcodes used to report on each biological feature probed.

### Novel Programs, Software, Algorithms

The CLR transformed datasets were analyzed through custom scripts in matlab2022b, with functions utilized from the Bioinformatics plug-in. Custom scripts are available on GitHub (https://github.com/mjd67/Multiple_RAM). Plots were produced using the library package(ggplot2) for R compiler x64 4.0.3 and written in Rstudio 2023.12.1 4.02.

### Web Sites/Data Base Referencing

The ViennaRNA package 2.0 via System2() in R compiler x64 4.0.3 and written in Rstudio 2023.12.1 4.02 was used for all RNA structure and folding predictions (32). QuantStudio™ 3/5 Real-Time PCR Design and Analysis Software v2 was used to analyze all qPCR data. Cq values were converted to copy numbers using standard curves generated from gel-purified amplicon DNA produced by PCR or RT-PCR with the corresponding primer set.

### Plasmid construction

Orthogonal barcodes were generated using vectors that transcribe cat-RNA-v1 (pPK076) and cat-RNA-v2 (pLNK007) using either Gibson assembly or Golden Gate cloning (33, 34). The promoter sequences used to transcribe cat-RNA are in Supplemental Table 2, while the cat-RNA barcode sequence variation is in Supplemental Table 3. Amplicons were generated using PCR reactions (50 µL) containing primers (0.5 µM each), template (≤1 ng), and one of two DNA polymerase mixtures: (i) 25 μL 2 × Phanta Max Master Mix, or (ii) 0.5 μL Q5® High-Fidelity DNA Polymerase, 25 μL of 5x Q5®Reaction Buffer, and 0.5 µM dNTPs. Golden gate reactions (15 µL) consisted of 25 fmol of each DNA product, 0.5 μL T4 DNA Ligase, 1.5 μL 10x T4 ligase buffer, and 0.5 μL of either BsaI-HFv2, BbsI-HF, or Esp3I. To generate plasmids for that transcribe inactive cat-RNA, the guanine in the catalytic core participates in a critical G·U wobble pair with the target rRNA was mutated to an adenine (18). All plasmids were sequence verified.

To create vectors that transcribe cat-RNA-v1 with orthogonal barcodes (Supplemental Figure 2), pPK076 was amplified to linearize the vector between the RT-qPCR probe and reverse transcriptase primer binding sites, yielding an amplicon flanked by Esp3I sites. This region was chosen to allow for the creation of sequence variation before the mRNA barcode that is amended to 16S rRNA. In addition, a pair of oligonucleotides, 13 base pairs each, containing the sequence NNNYRNNNYRNNN were synthesized by Integrated DNA Technologies (Coralville, Iowa). These oligonucleotides were mixed (2 µM) in a 25μL reaction that contained the primers, Q5® High-Fidelity DNA Polymerase, Q5®Reaction Buffer, and dNTPs. The reaction was incubated at 72°C for 20 minutes. The double stranded DNA (dsDNA) product of this reaction was cloned into the linear vector using Golden Gate cloning and transformed into *E. coli* MegaX using electroporation. This transformation was split evenly across 10 plates, which yielded ∼400 cfu/plate, which was estimated to represent 0.01% of the theoretical variants in the library. Individual colonies were used to inoculate cultures, and their plasmids were purified and sequenced using whole plasmid sequencing to determine barcode sequences and ensure proper golden gate assembly.

To create vectors that transcribe cat-RNA-v2 with orthogonal barcodes, pLNK007 was amplified using primers that eliminate the anti-Shine-Dalgarno anticodon middle stem-loop of the tRNA barcode, using an approach that mirrored pPK076 amplification. Pairs of complimentary oligonucleotides were synthesized that encoded each orthogonal barcode, which when annealed yield a dsDNA with single stranded overhangs. To anneal, 0.5 µL each oligonucleotide (100 µM) was mixed with 9 µL of water, heated to 95° in BioRad T100 thermocycler (Hercules, California), and ramped down to 37°C at a rate of 0.1°/sec. The product of this reaction was phosphorylated by mixing the 10 µL oligo mixture with 7.75 µL of water, 2 µL of NEB T4 DNA Ligase Reaction Buffer (B0202S) and 0.25 µL of NEB T4 polynucleotide kinase (M0201S). This solution is incubated at 37°C for >1 hr. They then were phosphorylated by mixing the annealed oligonucleotide (10 µL) with 2 µL of T4 DNA Ligase Reaction Buffer, 0.25 µL of T4 polynucleotide kinase, and 7.75 µL dH_2_O. This reaction was incubated at 37°C for 1 hr, cloned into the linearized cat-RNA-v2 using a two-part GoldenGate cloning, and transformed into *E. coli.* Individual colonies were used to inoculate cultures, their plasmids were purified, and whole plasmid sequencing was used to verify.

### Bacterial growth and transformations

*Escherichia coli* was made chemically competent by growing cells overnight in Lysogeny Broth (LB) at 37°C while shaking at 225 rpm, diluting 1:500, and growing the diluted cells to mid logarithmic phase. Cells were placed on ice for 10 min, harvested by centrifugation, and resuspended in LB containing 1% w/v PEG 3500, 0.5% v/v DMSO, and 20 mM MgCl_2_ such that cells were concentrated ∼40-fold. Prior to use, cells were stored at −80°C. Transformations were performed by mixing thawing cells (50 µL) with plasmids (1 ng), incubating on ice for 30 min, and heat shocking at 42°C for 30 seconds. After incubating on ice for 5 min, SOC medium (150 µL) was added, which contained 2% tryptone, 0.5% yeast extract, 10 mM NaCl, 2.5 mM KCl, 10 mM MgCl_2_, 10 mM MgSO_4_, and 20 mM Glucose. After incubating at 37°C while shaking at 225 rpm for 1 hour, cells were spread on LB-agar medium containing 50 µg/mL kanamycin and grown overnight at 37°C to obtain single colonies. All barcoding experiments for both strains of *E. coli* (DH10B and MegaX) were performed using cells grown in LB medium containing 50 µg/mL kanamycin at 37°C while shaking at 225 rpm.

*Vibrio natriegens* was made competent by growing in LB medium (50 mL) supplemented with V2 salts (204 mM NaCl, 4.2 mM KCl, 23.1 mM MgCl_2_) for 2 hrs at 30°C while shaking at 250 rpm. When cells reached mid to late logarithmic phase, they were pelleted by centrifugation, washed 3x times using electroporation buffer (680 mM sucrose and 7 mM K_2_HPO_4_, pH 7), resuspended in electroporation buffer (0.5 mL), and frozen in aliquots (50 µL) at −80°C. To transform, an aliquot was mixed with plasmid DNA (100 ng), incubated on ice for 1 min, and electroporated using a 0.9 kV pulse for ∼5 ms in a 0.1 cm cuvette. After recovery in LB (0.5 mL) containing 680 mM sucrose and V2 salts for 1 hr at 30°C while shaking at 250 rpm, cells were plated on LB-agar plates supplemented with V2 salts and kanamycin (200 µg/mL) and grown overnight at 30°C to obtain single colonies. All barcoding experiments were performed using cells grown in LB-v2 medium containing kanamycin (200 µg/mL) at 30°C while shaking at 225 rpm.

Electrocompetent *P. putida* were prepared by culturing cells in LB medium (50 mL) at 30°C overnight while shaking at 225 rpm in a 250 mL flask. Cells were pelleted by centrifugation, washed with 10% glycerol (25 mL) three times, and resuspended in 10% glycerol (1 mL). Competent cells were either stored in 50 µL at −80°C or used immediately for electroporation. For transformations, competent cells were mixed with plasmid DNA (100 ng), incubated for 30 minutes at 25°C, and electroporated using a 1.6 kV pulse for ∼5 ms with a 0.1 cm cuvette. After adding SOC medium (1 mL), cells were incubated at 30°C for 1 hr while shaking at 225 rpm, cells were plated on LB-agar medium containing kanamycin (50 µg/mL), and plates were incubated overnight at 30°C to obtain single colonies. All barcoding experiments were performed using cells grown in LB medium containing 50 µg/mL kanamycin at 30°C while shaking at 225 rpm.

### Barcoding experiments

The 16S rRNA sequences used to differentiate microbes are provided in Supplemental Table 4. To evaluate the barcoded-rRNA signal of orthogonal cat-RNA transcribed from the same promoter, each strain (*Ec*:*Vn*:*Pp*) was transformed with individual plasmids that transcribe the different orthogonal cat-RNA. These microbes were analyzed using RT-qPCR. Prior to NGS, equivalent titers of each transformed strain were mixed to create single strain cultures with multiple orthogonal cat-RNA. These single-strain cultures were mixed at stoichiometries of 1:19:80 (*Ec*:*Vn*:*Pp*) prior to assessing barcoding signals in a community. To evaluate barcoded-rRNA from orthogonal cat-RNA-v2 transcribed from four different promoters in plasmids having two different origins of replication, *E. coli* was transformed with the eight different plasmids individually, which were characterized using RT-qPCR, and then mixed at equivalent titers prior to NGS. To evaluate barcoding from these eight plasmids across three microbes, five orthogonal cat-RNA were used to report on each promoter and origin combination (n = 40 plasmids). Plasmids having different promoters but the same origin of replication were mixed at equivalent stoichiometries, yielding two plasmid pools, and these were transformed into *E. coli* using heat shock, conjugated into *V. natriegens*, and electroporated into *P. putida*. Only pBBR1 origins were transfected for *P. putida* due to incompatibility with pColE1 oriV. The transformants were harvested from plates, and the cell slurries were used to inoculate LB (*E. coli and P. putida*) or LBV2 (*V. natriegens*) cultures containing kanamycin (50 µg/mL) at an OD of 0.01, which were grown to ODs of ∼0.4 (*E. coli*), 0.5 (*P. putida*), and 0.25 (*V. natriegens*). RNA was extracted from each culture and mixed ratios 6:2:1 (*Ec:Vn:Pp*) to produce an even representation of the three species following NGS. All samples were split into three aliquots to allow for: (i) RNA extraction, which yielded total 16S rRNA and barcoded-rRNA for NGS, (ii) plasmid extraction, which enable us to quantify the relative abundances of the different plasmids in each sample, and (iii) genomic DNA extractions, which allowed for analysis of the amount of each organism in the mixtures. Following mixing, cells in each aliquot were pelleted, the supernatant was removed, and pellets were stored at −80°C until analysis.

### RNA extraction

A modified version of the MagMAX™ Microbiome Ultra Nucleic Acid Isolation Kit was used with a KingFisher Apex System. In brief, each pellet was resuspended in MagMAX™ Microbiome Lysis Solution (400 µL) and treated with MagMAX™ Viral/Pathogen Proteinase K (40 µL). Prior to the elution step, a mixture of TURBO DNAse (10 µL of 10 U/mL) and buffer (190 µL) were added to each sample and incubated for 30 minutes at 37°C. Samples were then mixed with nucleic acid binding beads, which were washed prior to elution using MagMAX™ Viral/Pathogen Elution Buffer (50 µL).

### DNA extraction

Plasmid DNA was extracted from microbes using the QIAprep Spin Miniprep Kit, and the purified DNA was used as a template for generating amplicons prior to NGS. Genomic DNA was extracted from synthetic communities using the Wizard® Genomic DNA Purification Kit, and a fragment of the 16S rRNA gene was amplified prior to NGS.

### Reverse transcription quantitative PCR

Different combinations of primer and probe sequences (Supplemental Table 5) were used to quantify: (i) native 16S rRNA, (ii) barcoded-rRNA generated by cat-RNA-v1 and cat-RNA-v2, (iii) total cat-RNA by targeting the catalytic region of the ribozyme, and (iv) total cat-RNA barcode, which represents the sum of the spliced and unspliced barcode. To quantify native 16S rRNA, RT-qPCR reactions were performed using a dye-based qPCRbio Sygreen 1-step kit (PCR Biosystems, PB25.11). RNA extractions were diluted 10,000x for native 16S rRNA analysis. Reactions (20 µL) consisted of 0.4 μM of each primer, 1 µL of PCRbiosystems 20x RTase Go reverse transcriptase with RNAse inhibitors (PB10.53-05), and 8 μL of diluted RNA extract as template. To quantify cat-RNA, barcoded-rRNA, and total barcode, RT-qPCR reactions were performed with probes using TaqMan™ Fast Virus 1-Step Master Mix (Applied Biosystems). With each reaction (20 µL), 100x diluted RNA extraction (8 μL) was added with 0.9 μM of each primer and 0.25 μM of each probe. RNA-free water was used to reach the final volume.

### Next generation sequencing

Four types of amplicons were generated using different primer sets (Supplemental Table 5). First, to generate 16S rRNA amplicons (∼450 bp), RT-PCR was performed using LunaScript® Multiplex One-Step RT-PCR Kit in a Biorad T100 thermocycler. Each reaction (25 µL) contained 1.25 μL DMSO, 5 μL of reaction buffer, 1 μL of enzyme mix, 0.5 μM of each native 16S RT-NGS primer (oRM20 and oRM18), and 4 μL of RNA template which was a 100x dilution of the extracted RNA. Second, to produce barcoded-rRNA amplicons arising from cat-RNA-v1 (480 bp) and cat-RNA-v2 (515 bp), the same protocol was used with primers that amplify barcoded-rRNA containing cat-RNA-v1 (oRM18 and oRM21) and cat-RNA-v2 (oRM18 and oMJD154) barcodes. Third, to generate amplicons of cat-RNA-v2 (280 bp) from plasmid extracts, PCR (25 µL) was performed using 0.5 μM primers (oMJD99 and oMJD154), template (≤1 ng), 0.25 μL of Q5® High-Fidelity DNA Polymerase, 25 μL Q5®Reaction Buffer, and 0.5 μM dNTPs. Fourth, to generate amplicons of 16S rRNA (450 bp) from genomic DNA extracts, PCR (25 µL) was performed using the same protocol with native 16S rRNA NGS primers. In all cases, amplicons identified by agarose gel electrophoresis were purified before sequencing. Amplicons were sequenced by Plasmidsaurus (South San Francisco, CA) using Oxford Nanopore Long-read NGS or by Genewiz (South Plainfield, NJ) using Illumina Short-read NGS.

### RNA stability calculations

For both cat-RNA v1 and v2, calculations were performed using ViennaRNA (32). For these calculations, we used the entire barcode sequence fused to the last 20 bp of the ribozyme as the input sequence. For barcoded-rRNA predictions, we used the entire barcode sequence and 20 bp upstream of the targeted *E. coli* 16S rRNA splice site. Minimal free energy (MFE), optimal secondary structure, and positional entropy were calculated using RNAfold with parameters -p and --noPS. The base pair distance of barcode variants to the original design template was calculated using RNAdistance with parameters -Xp, -D, and P. Rationally designed barcodes were generated using RNAinverse with the default parameters using the RNAfold-predicted secondary structure of the original tRNA design as the input structure. In total, 40 rationally designed barcodes that had the optimal local entropy at the stem flanking the internal barcode-containing loop structure were selected for testing in cells. MFE, positional entropy, and base pair deviation from the original cat-RNA design for each internal barcode variant are reported in Supplemental Table 6.

### Data analysis pipeline

Supplemental Figure 3 summarizes the custom pipeline used for short read amplicon NGS data analysis. Four NGS data sets were analyzed, including: (i) 16S rRNA amplicons generated from RNA extracts, (ii) barcoded-rRNA amplicons from RNA extracts, (iii) cat-RNA-v2 barcodes from plasmid extracts, and (iv) 16S rRNA amplicons from genomic DNA. Briefly, all raw files isolated FASTQ quality scores for filtering with built in function *FASTQread*(). Primers were trimmed off and incomplete reads by total length were removed. For (i,ii and iv), species identification was implemented. Barcoded cat-RNA-v2 paired reads lack overlap and required a gap for alignments. 16S rRNA sequences used for alignment are in Supplemental Table 4 and used function *nwalign*(). For (ii and iii) barcode identification was implemented. Reverse reads were trimmed to isolate barcodes and only exact matches were accepted as a confirmed read. For (ii) CLR transformations were computed on pooled bins by species and construct based on sequence references as shown in Supplemental Table 3. All CLR data processing involved removed pooled elements which had any value less than the limit of detection (LOD = 10 reads), in any of the sequencing runs. Logarithms cannot be performed on values equal to zero, a requirement of compositional data analysis, and as such must be either removed or set to a percentage of the LOD (35, 36). Data processing and formatting differed for nanopore datasets. Firstly, Cutadapt was used to trim off forward and reverse primers, in both orientations, and discard untrimmed reads with default parameters. After primer trimming, reads were screened for the expected universal barcode sequence (5’ end of the barcode) in both forward and reverse orientations. Data was then imported into QIIME 2 (37). Briefly, DADA2 were used to filter, demultiplex, remove chimera, and generate ASV sequences and feature tables. Taxonomy was assigned to each ASV sequence using *feature-classifier classify-sklearn* with the preformatted SILVA 138 SSURef NR99 full-length classifier.

## RESULTS

### Orthogonal cat-RNA-v1 present high signal variability

The prototype RAM system uses a cat-RNA that splices a barcode derived from a fragment of the *gfp* transcript onto 16S rRNA in between variable regions V8 and V9 (18), referred to herein as cat-RNA-v1 (Figure 1). Previously, a single orthogonal cat-RNA-v1 was created by inserting a five base pair sequence into the barcode, and direct comparison of the two orthogonal cat-RNA-v1 signals in *E. coli* yielded similar barcoded-rRNA levels (18), suggesting that small sequence changes in the barcode can be used in cat-RNA-v1 to create an orthogonal barcoding system. To evaluate if this design approach can be used to create additional orthogonal cat-RNA that perform consistently, thirteen random nucleotides were inserted in cat-RNA-v1 to yield a library of vectors. These vectors were modified to transcribe orthogonal cat-RNA-v1 under the control of 23 different synthetic promoters, with each promoter represented by three unique barcodes. Following transformation into *E. coli*, RT-qPCR was used to evaluate the signals for the trios of vectors representing each promoter. Across all promoters, the cells transcribing cat-RNA-v1 presented barcoded-rRNA signals that varied pairwise by up to >100-fold (Supplemental Figure 4A). Among the orthogonal cat-RNA-v1 transcribed from the same promoter, the average variation across all trios was 9.2-fold. The lowest variation observed for a trio was 1.9-fold, while the greatest variation was 75.2-fold. When NGS was used to analyze the same library (Supplemental Figure 4B), high variation was also observed. The raw count totals were linearly transformed by the centered log-ratio (CLR), under the assumption that trios of barcodes function as biological replicates. This CLR transformed data represents the relative abundances of each barcode across the complete dataset, with zero representing the geometric average abundance of all barcodes. In line with the RT-qPCR results, the CLR transformed sequencing counts also showed high average internal variation among all cat-RNA-v1 designs (15.2-fold), with a range from 2.4-fold to 70.3-fold across trios sharing the same promoter. These results show that barcode sequence variations within cat-RNA-v1 produce high signal variability across orthogonal cat-RNA transcribed from the same promoter.

To better understand the effect of cat-RNA-v1 sequence variation on barcoded-rRNA levels, twelve orthogonal cat-RNA-v1 with twelve unique barcodes were used to report on the same synthetic promoter (P1) in a plasmid with a pBBR1 origin of replication (Figures 2A-B); these barcodes were chosen from a large library of randomly-generated barcodes. RT-qPCR of the barcoded-rRNA in *E. coli* revealed that the highest and lowest signals varied by 13.5-fold (Figure 2C), with an average pairwise variation of 5.4-fold. To ascertain if the variability arose due to low transcriptional rates, a stronger synthetic promoter (P18) was used to drive the transcription of the same twelve orthogonal cat-RNA-v1. With P18, the highest and lowest signals varied by 4.7-fold, with a smaller average pairwise variation of 2.7-fold. A comparison of the data from each promoter revealed that while the number of copies of each barcoded-rRNA increased in *E. coli* with P18 (Figure 2E), a strong correlation (R^2^ = 0.75) was observed between the barcoded-rRNA signals for the same orthogonal cat-RNA-v1 transcribed by promoters P1 and P18. Analysis of variation across each set of promoter data using a Kruskal-Wallis rank order test revealed significant variation in signals arising from orthogonal cat-RNA-v1 transcribed by promoter P1 (n = 12; p = 0.01) and P18 (n = 12; p = 0.002). Together, these results show that the sequence changes within the cat-RNA-v1 barcode is a major contributor to the signal variation observed with barcoded-rRNA in *E. coli*.

**Figure 2.**
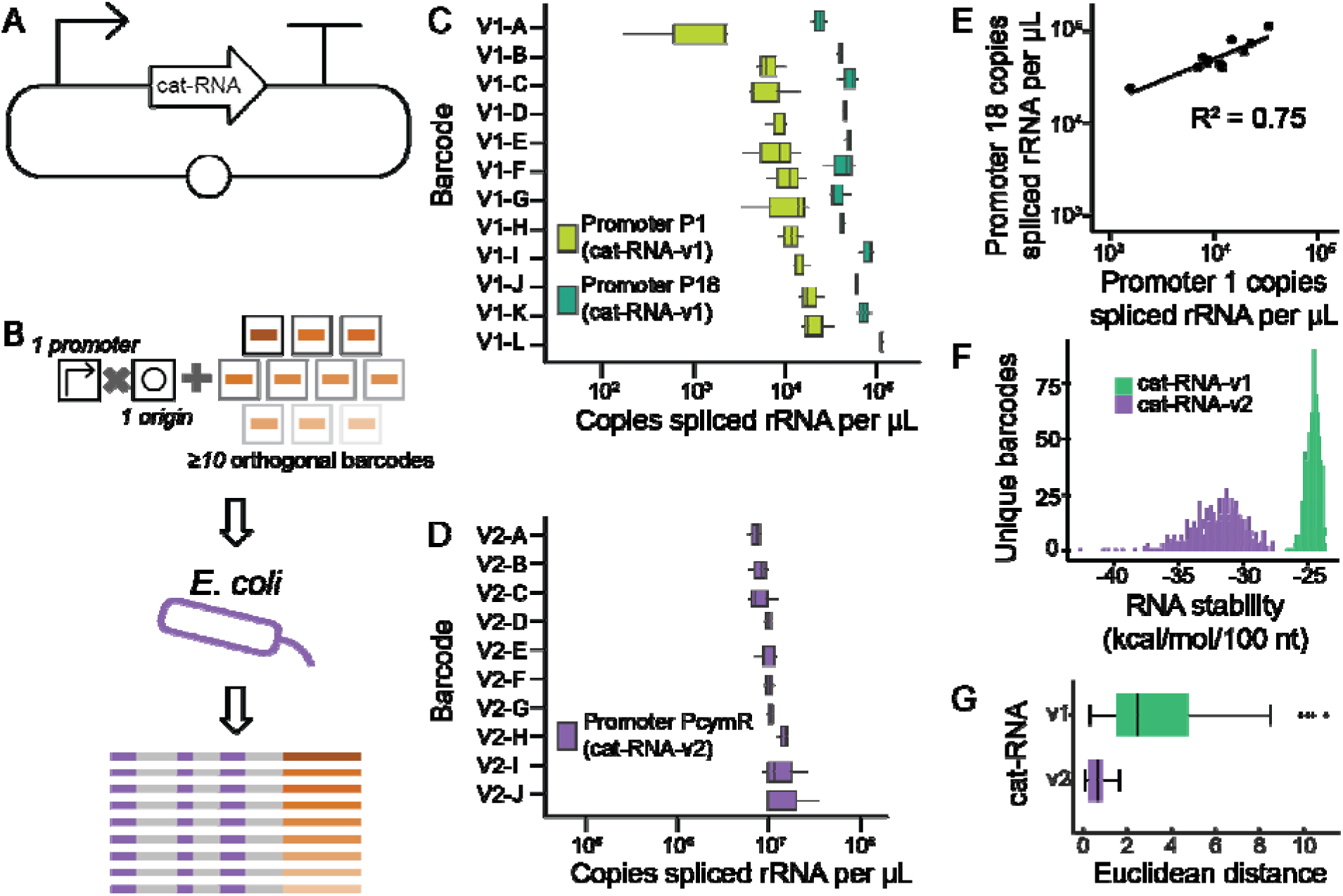
Sequence modifications in the cat-RNA barcode affect the RAM signal. **(A)** The plasmids used to compare orthogonal cat-RNA. (**B**) All plasmids used the same promoter to transcribe cat-RNA with orthogonal barcodes and contained a pBBR1 origin of replication. (**C**) Variability in the RT-qPCR signal (3 biological replicates) of barcoded-rRNA arising from orthogonal cat-RNA-v1 transcribed using the P1 (light green) and P18 (dark green) promoters. (**D**) RT-qPCR signal (3 biological replicates) of barcoded-rRNA arising from orthogonal cat-RNA-v2 transcribed by P_cym_. (**E**) Orthogonal cat-RNA-v1 transcribed using P1 and P18 exhibit a linear correlation (R^2^ = 0.75). **(F)** Stability modeling for 500 orthogonal cat-RNA-v1 (green) and cat-RNA-v2 (purple) barcodes. **(G)** Euclidean distances of orthogonal barcodes derived from CLR transformed counts of NGS data. The cat-RNA-v1 distances were calculated using the trios of orthogonal barcode signal differences observed across 23 unique promoters (# of distances = 69), across three biological replicates (n = 69 x 3 = 207), while the cat-RNA-v2 distances used pairwise comparisons of 10 orthogonal barcodes, with 45 pairwise distances, across five biological replicates (n = 5 x 45 = 225). The box and whisker plots represent the full set of pairwise distances between all orthogonal barcodes, which is a representation of how much influence barcode sequences have on barcoded-rRNA levels.

To investigate why sequence changes in the cat-RNA-v1 barcode led to signal variability, the total barcode was quantified using RT-qPCR, which represents the sum of the barcoded-rRNA and the unspliced cat-RNA-v1. When total barcode concentration was evaluated in *E. coli* transcribing the same set of twelve orthogonal cat-RNA-v1 barcodes using the P1 and P18 promoters (Supplemental Figure 5), the correlation across promoters was weaker (R^2^ = 0.37) than that observed for barcoded-rRNA alone. Since total barcode abundance was previously found to represent a proxy for promoter activity (18), this weaker correlation suggests that barcode sequence variation differentially affects the stability of both unspliced cat-RNA-v1 and the barcoded-rRNA product.

To evaluate whether barcode sequence changes affect the stability of cat-RNA-v1 prior to trans-splicing, an inactivating mutation (G264A) was incorporated into six of the orthogonal cat-RNA-v1 barcodes in constructs containing promoters P1 and P18, designated d.cat-RNA-v1. Previously, this mutation was found to inactivate cat-RNA-v1 without altering steady-state levels (38). When RT-qPCR was used to evaluate orthogonal cat-RNA-v1 abundances (Supplemental Figure 6), the total cat-RNA-v1 signals differed by 1.7-fold when transcribed from the P1 promoter, while a 2.5-fold difference was observed with the P18 promoter. In addition, the rank order was no longer correlated (R^2^ = −0.09) across the two promoters. Further, the rank order for the barcoded-rRNA signals arising from active cat-RNA-v1 with different barcodes was distinct from the rank order of those with inactive cat-RNA-v1 (Supplemental Figure 7). These trends suggest that the large variance with orthogonal cat-RNA-v1 arises from changes in the stability of barcoded-rRNA products rather than changes in the steady-state transcript levels of orthogonal cat-RNA-v1.

### Structured cat-RNA-v2 barcodes present lower inter-barcode variability

A recent study showed that cat-RNA-v2 improves barcoded-rRNA signal by more than an order of magnitude by using a structured barcode derived from *E. coli* transfer RNA (tRNA) (31). We hypothesized that the increased secondary structure of the cat-RNA-v2 barcode, specifically the anti-Shine-Dalgarno anticodon stem–loop region, would limit the impact of orthogonal barcode sequences on barcoded-rRNA signal variability, since this region has evolved to contain sequence diversity in nature (39). To compare the stability of orthogonal barcodes in cat-RNA-v1 and cat-RNA-v2, we used the RNA folding prediction algorithm ViennaRNA to estimate the thermodynamic stability of 500 orthogonal barcodes generated within each of the cat-RNA-v1 and cat-RNA-v2 designs (32). Orthogonal cat-RNA-v1 barcodes presented more positive adjusted minimum free energies (AMFE) than orthogonal cat-RNA-v2 designs (Figure 2F), indicating that sequence variation within cat-RNA-v1 produces less stable barcode structures than equivalent variation within cat-RNA-v2.

To directly measure the effects of sequence variation on cat-RNA-v2, we rationally varied 10 bp in the anti-Shine Dalgarno anticodon stem–loop. Using AMFE predictions,10 orthogonal barcodes were selected that display the same predicted structure as the original cat-RNA-v2 and a minimum free energy folding prediction within 1 unit kJ/mol per 100 nucleotides (Supplemental Table 6). Vectors transcribing these orthogonal cat-RNA-v2 were transformed into *E. coli*, and the barcoded-rRNA signal from each was measured using RT-qPCR (Figure 2D). All ten cat-RNA-v2 presented >10-fold higher barcoding signals compared to the orthogonal cat-RNA-v1. In addition, the maximum pairwise difference in signal between any two barcodes was 2.5-fold, and no significant difference in barcoded-rRNA was observed across the ten orthogonal cat-RNA-v2 (Kruskal-Wallis test; p = 0.13). These findings show that orthogonal barcodes inserted within the anti-Shine Dalgarno anticodon stem-loop of cat-RNA-v2 produce significantly lower inter-barcode signal variability than equivalent sequence variation in cat-RNA-v1.

When RAM is used in communities, the signal is read out using next generation sequencing (NGS) of barcoded-rRNA (18, 19). To assess whether cat-RNA-v2 presents a more consistent signal when detected with NGS, we compared the relative abundances of barcoded-rRNA in cultures containing similar titers of cells containing orthogonal cat-RNA-v1 or cat-RNA-v2. For cat-RNA-v1, *E. coli* was transformed with the 69 constructs representing 23 trios of orthogonal cat-RNA-v1 barcodes. To create a cell mixture containing equal proportions of each orthogonal cat-RNA-v1, cells transformed with each construct were grown to mid-exponential phase and mixed at equal titers. Three subsamples were taken from which total RNA was extracted. For cat-RNA-v2, a mixture of *E. coli* transformed with the ten orthogonal variants was prepared in a similar manner, except for five subsamples were taken from which total RNA was extracted. Total RNA was extracted from these mixtures and subjected to one-step RT-PCR to produce barcoded-rRNA amplicons, which were gel purified and sequenced. The resulting amplicon sequencing data was linearly transformed using a CLR transformation (Supplemental Figure 2A-C) because: (i) the distances between barcoding signals, and from the geometric mean, are conserved with this transformation, and (ii) the transformed data occupy Euclidean space, enabling the application of traditional statistics such as t-tests (Supplemental Figure 2D). We compared the Euclidean distances between all orthogonal barcodes for each cat-RNA design (Figure 2G); for this data, a distance of zero represents a perfect design in which no difference between orthogonal barcodes is observed. The cat-RNA-v1 designs had a mean distance of 3.29, which was five times greater than that of the cat-RNA-v2 designs (0.65). When correcting for the log_2_ scale of the Euclidean distances, orthogonal cat-RNA-v2 exhibited 32-fold lower inter-barcode variability than orthogonal cat-RNA-v1. Thus, these results demonstrate that orthogonal cat-RNA-v2 designs present significantly lower variability than cat-RNA-v1 designs, indicating that cat-RNA-v2 is better suited for multiplexed RAM experiments.

### Estimating the number of orthogonal cat-RNA to resolve biological features

Given the inter-barcode variation observed in barcoded-RNA signals from orthogonal cat-RNA transcribed using the same promoter, we sought to determine how the number of orthogonal barcodes used as replicates affects the ability to detect differences in RAM signal of a given magnitude (i.e., the ability to use cat-RNA-v1 or cat-RNA-v2 to report on different biological processes in parallel, such as the activities of different promoters). To do this, we applied a power analysis framework commonly used to determine the minimum effect size detectable given a defined number of replicates and an empirical noise estimate (38). The Euclidean distances of orthogonal cat-RNA-v1 and cat-RNA-v2 splice rRNA signals were fit to a half-normal distribution using a generalized linear model, providing an empirical noise estimate for each design. By dividing the sum of squares between (SSB) barcode distances by the square root of the number of orthogonal barcodes (*n*), this equation predicts the minimum fold-change in barcoded-rRNA signal that can be distinguished from background noise, *i.e.*, the limit of detection, as a function of barcode replicate number.

The relationship between the number of orthogonal barcodes required per library element (i.e., biological feature) and the detectable fold changes is logarithmic, with diminishing returns as barcode number increases and the curve asymptotes towards the inherent error of the system. When applied to the cat-RNA-v1 library in *E. coli* (Supplemental Figure 4), which contained 3 replicates of 69 Euclidean distances for total of 207 unique distances, we found that 5 barcodes would be required per library element to observe a 5-fold difference in barcoded-rRNA signal (Figure 3A). In other words, to detect a 5-fold difference across two different promoters in a pure *E. coli* culture using NGS analysis of cat-RNA-v1 barcoded-rRNA, each promoter would require 5 orthogonal cat-RNA-v1. By contrast, when the same analysis was applied to the cat-RNA-v2 dataset for *E. coli* (Figure 2D), which contained 5 replicates of 45 Euclidean distances for a total of 225 unique distances, only a single orthogonal barcode is required per library element to resolve a 5-fold difference in signal in a pure *E. coli* culture using NGS. These results provide a quantitative framework for evaluating how the lower inter-barcode variability of cat-RNA-v2 decreases the number of orthogonal barcodes required to resolve biological features. Due to the lower variability of orthogonal cat-RNA-v2, this design was used for all subsequent experiments.

**Figure 3.**
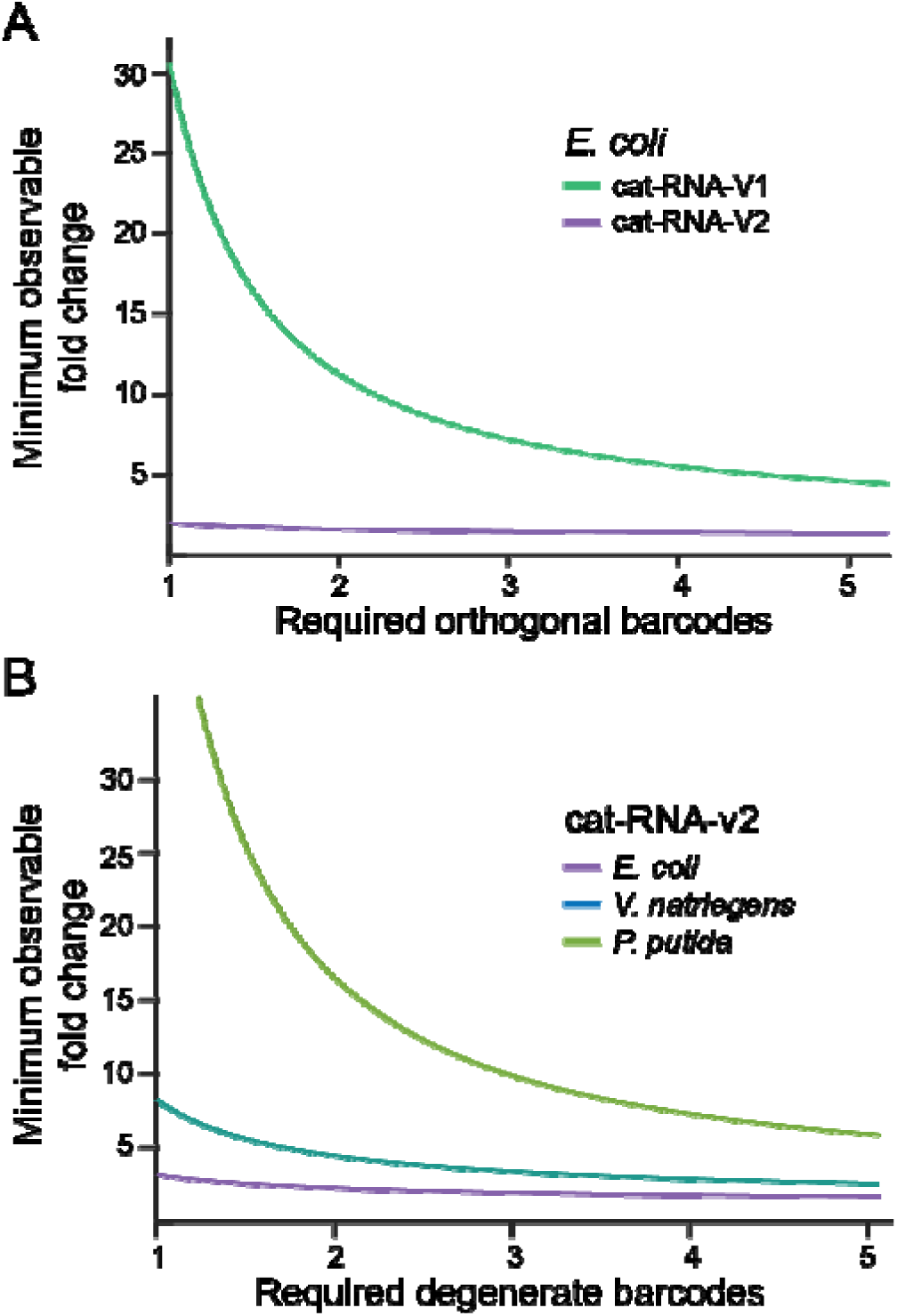
Estimate of the number of degenerate cat-RNA needed to resolve signal differences across plasmids using NGS. The sum of squares of the Euclidean distances between the orthogonal barcodes within an organism was divided by square root of each number of degenerate barcodes per element to predict observable fold change of a given composition. Using experimental measurements of orthogonal cat-RNA, we used this relationship to estimate the number of barcodes required to observe a specific fold change: (**A**) within *E. coli* using either cat-RNA-v1 or cat-RNA-v2 to compare biological features, and (**B**) across organisms using cat-RNA-v2 to compare biological features.

### Multiplexed barcoding by ten orthogonal cat-RNA-v2 within a microbe

To test if we could multiplex information storage across different organisms, we evaluated signals of the ten orthogonal cat-RNA-v2 barcodes in *E. coli*, *Vibrio natriegens*, and *Pseudomonas putida* (Figure 4A). We first measured the concentration of barcoded-rRNA produced by each orthogonal design individually in each organism using RT-qPCR and normalized these signals to total 16S rRNA (Figure 4B). *E. coli* presented the highest signals, while *P. putida* presented the lowest signals, and *V. natriegens* had intermediate signals. Significant inter-barcode variation was observed in *P. putida* (Kruskal-Wallis; p = 0.02), while no significant variation was observed in *E. coli* (p = 0.4) or *V. natriegens* (p = 0.2). These results show that the signal variability observed with orthogonal cat-RNA-v2 is inversely proportional to cat-RNA expression.

**Figure 4.**
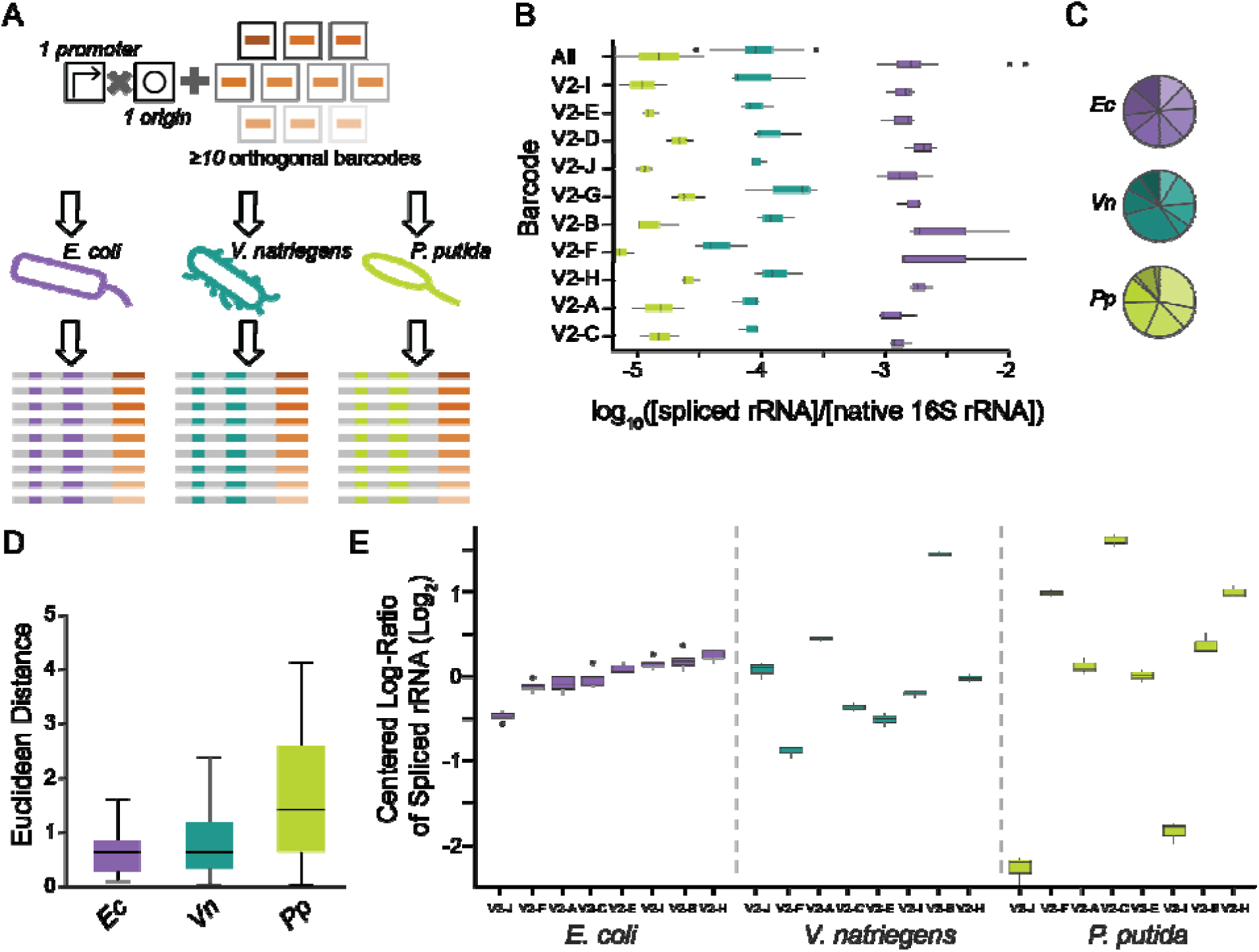
Signal variation for eight orthogonal cat-RNA-v2 across three microbes. **(A)** Plasmids with a pBBR1 replication origin were created that transcribe eight orthogonal cat-RNA-v2 using P_cym_. Following transformation into *E. coli* (purple)*, V. natriegens* (teal), and *P. putida* (green), the barcoded-rRNA signal was monitored in (**B**) isolates using RT-qPCR using 3 biological replicates per strain and (**C**) cell mixtures using NGS with 3 sequencing replicates. The RT-qPCR barcoded-rRNA signal was normalized to total 16S rRNA, and the average from all ten barcodes (all) was calculated (n = 30). **(D)** The Euclidean distances, derived from all pairwise distances between the CLR transformed counts of each pair of barcodes for each organism (n = 3 sequencing replicates, m = 28 unique pairwise distances). **(E)** CLRs were calculated using NGS for comparison, with barcodes ordered by their relative abundances in the *E. coli* sequencing data (n=3). The CLR transformed data represents the relative counts for each barcode in reference to the central tendency of the compositional sequencing dataset for each strain. Values above zero represent more abundant sequences, and those below zero are less abundant.

We next used NGS to read out the orthogonal cat-RNA-v2 signals from single-species cultures containing eight of the previously tested ten orthogonal designs. Two barcodes were excluded based on transformation success, to reduce the sample space to a single byte of information. Each strain was grown independently to mid-exponential phase, and individual strains containing each orthogonal barcode were pooled at equivalent titers prior to RNA extraction. In each pool, all eight barcodes were detected (Figure 4C). The Euclidean distances in the barcoded-rRNA reads (Figure 4D) were significantly different across all pairwise strain comparisons (Wilcox rank-sum; p < 0.005), with significantly greater inter-barcode variability observed in *V. natriegens* and *P. putida* than in *E. coli*. In addition, the rank order of signals across orthogonal barcodes varied across organisms (Figure 4E). These findings show that NGS can be used to read out barcoded-rRNA generated by eight orthogonal cat-RNA-v2 in parallel, and that there are species-specific influences upon barcoded-rRNA stability.

### Multiplexed barcoding by eight orthogonal cat-RNA-v2 within a community

To investigate whether orthogonal cat-RNA-v2 could be multiplexed in a synthetic community, we mixed *E. coli, V. natriegens,* and *P. putida* that had been transformed with the same eight orthogonal cat-RNA-v2 at a ratio of 1:19:80 (*Ec:Vn:Pp*) to account for differences in splicing efficiency across strains Following mixing, total RNA was extracted, barcoded-rRNA and 16S rRNA amplicons were generated, and NGS was used to quantify the species-level signals (Figure 5A). While the relative abundances of counts from each microbe varied (Supplemental Figure 8), all twenty-four barcodes were detected in the barcoded-rRNA pool, albeit at different abundances (Figure 5B-C). However, a comparison of the rank order of signal abundances across microbes revealed variation (Supplemental Figure 9). These results show that orthogonal cat-RNA-v2 barcodes can be used to simultaneously record and resolve host taxonomic information within a synthetic community.

**Figure 5.**
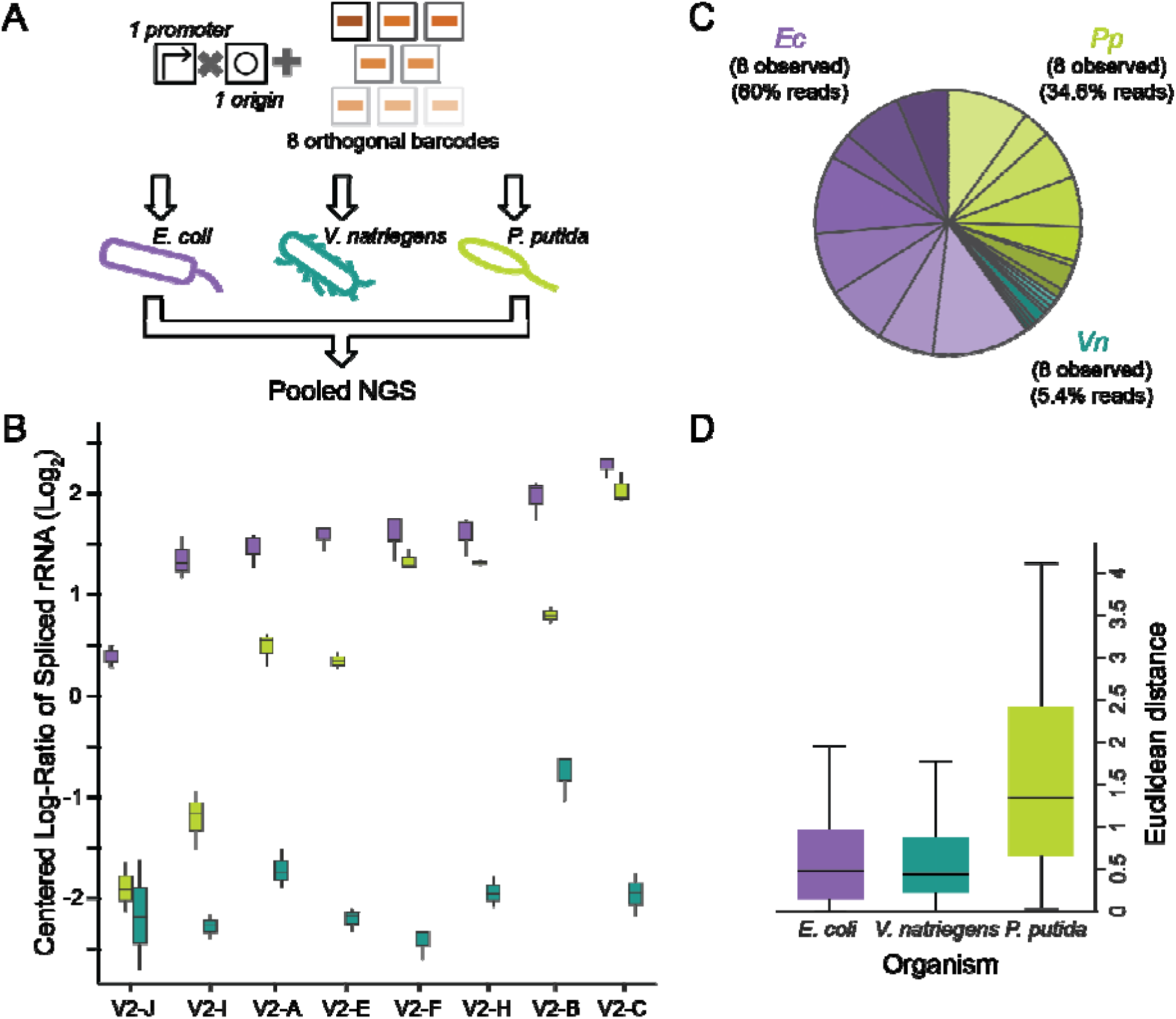
Abundances of eight orthogonal barcodes in a synthetic community. (**A**) The abundances of eight orthogonal cat-RNA were evaluated across a culture containing *E. coli* (purple)*, V. natriegens* (teal), and *P. putida* (green) transformed with the same eight plasmids that transcribe orthogonal cat-RNA-v2 from P_cymR_. (**B**) NGS analysis of barcoded-rRNA abundances in the mixed culture (3 sequencing replicates) revealed variation in the relative abundance of each barcode. (**C**) The average abundances of barcoded-rRNA signals varied across organisms. (**D**) The Euclidean distances, derived from the pairwise distance between the CLR transformed counts of each barcode to every other barcode within a replicate, for each organism in the mixed community (n = 3 replicates, m = 28 unique pairwise distances).

To evaluate how barcode sequences impact inter-barcode variability on a per-species basis within the community, we compared the Euclidean distances between orthogonal barcodes for each organism using barcoded-rRNA data from the synthetic community. *E. coli* had the lowest mean Euclidean distance (2.19), followed by *V. natriegens* (2.48), while *P. putida* had a significantly larger distance than the two other microbes (5.05; Wilcox rank-sum; p < 0.05) (Figure 5D). To understand the impact of using a mixture of microbes relative to monocultures, we compared the Euclidean distances from the synthetic community to those from the corresponding monocultures for each organism containing the same eight orthogonal cat-RNA-v2. *E. coli* exhibited significantly greater inter-barcode variability in the community compared to the monoculture (upper gate values for monoculture = 0.95, synthetic community = 2.19; p << 0.05), while *V. natriegens* showed a significant decrease in variability in the community relative to monoculture (upper gate values for monoculture = 2.48, synthetic community = 1.83; p = 0.016). No significance difference was observed for *P. putida* (upper gate values for monoculture = 5.48, synthetic community = 5.05; p = 0.88) (Supplemental Figure 10). These results show that community context influences inter-barcode variability across species, and that monoculture characterization of orthogonal cat-RNA-v2 cannot fully predict behavior in a mixed community.

Using the mathematical framework previously applied to evaluate cat-RNA-v1 and cat-RNA-v2, we apply our predictive model to evaluate cat-RNA v2 in other species. To do this, we fit the Euclidean distances of orthogonal cat-RNA-v2 barcoded-rRNA signals derived from the mixed communities, for each organism, to a half-normal distribution using a generalized linear model, providing an empirical noise estimate for each organism. All three organism data sets are composed of 28 distances across 3 replicates, for a total of 84 distances per organism. For both *E. coli* and *V. natriegens*, 5 orthogonal cat-RNA barcodes are sufficient to observe a 5-fold change between constructs in each species. However, *P. putida* does not reduce noise sufficiently, so 5 orthogonal cat-RNA-v2 barcodes are insufficient to detect a 5-fold change (Figure 3B). These results show that the lower inter-barcode variability of cat-RNA-v2 is not conserved across bacteria, and subsequent investigations are required to reduce noise outside *E. coli*.

### Storing 1 byte of information in rRNA using one cat-RNA-v2 per feature

We next sought to investigate whether RAM could report on eight biological features in a single strain using one orthogonal cat-RNA-v2 per feature. To do this, constructs containing two different origins of replication (pBBR1 and a ColE1) were built that use four different synthetic promoters to express eight orthogonal cat-RNA-v2 (Figure 6A). The promoters were chosen to span a range of predicted strengths (40), while the origins were selected because they differ in copy number, from ∼5 copies (pBBR1) to ∼15-50 copies (ColE1) (41–44).

**Figure 6.**
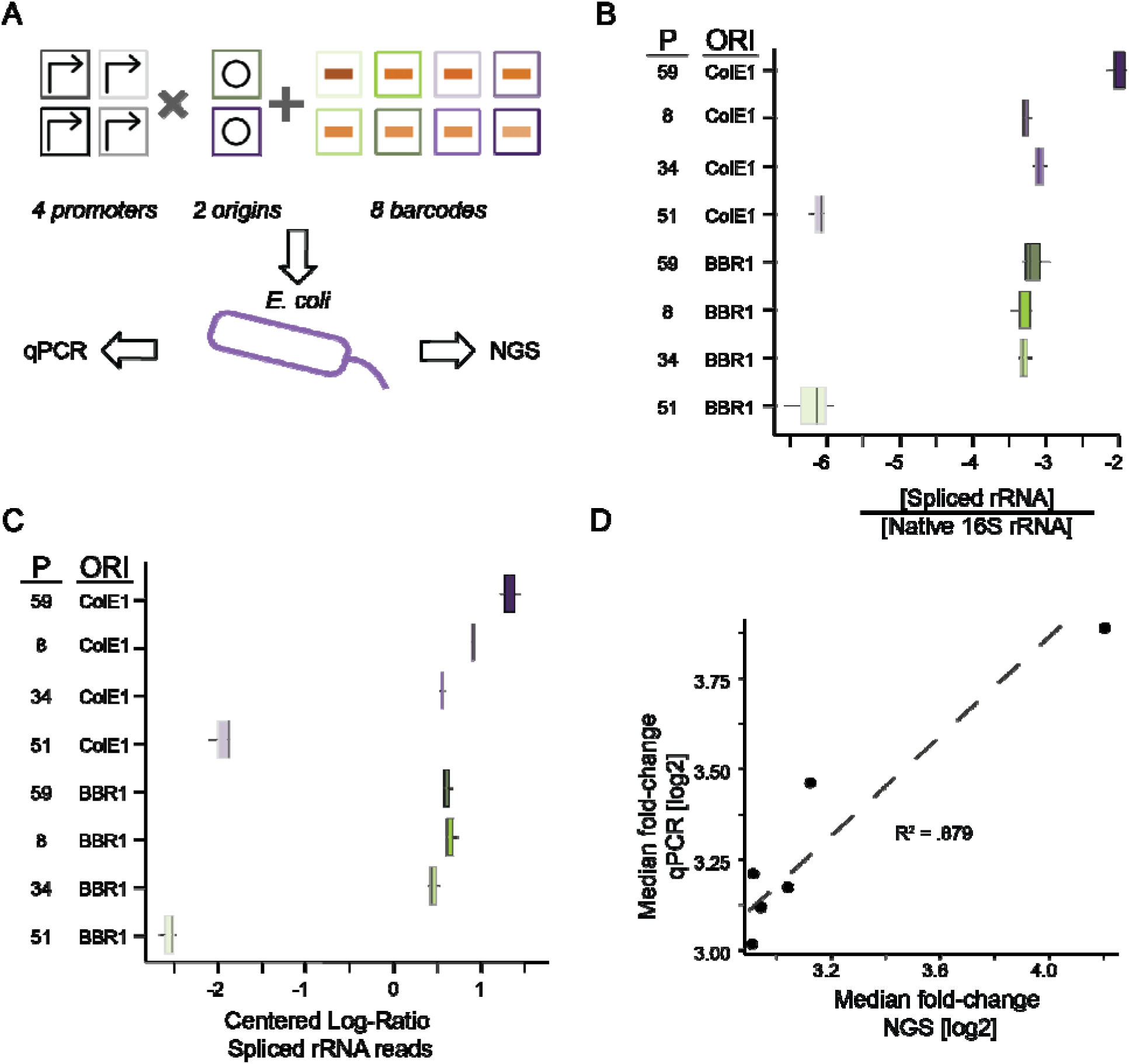
Recording 1 byte of plasmid information in *E. coli*. **(A)** Eight orthogonal cat-RNA-v2 were used to report on vectors that use different combinations of promoters (P8, P34, P51, and P59) and origins of replication (ColE1 and pBBR1). **(B)** RT-qPCR analysis of the barcoded-rRNA normalized to total 16S rRNA (3 biological replicates) from individual strains. (**C**) NGS analysis of barcoded-rRNA abundances from an *E. coli* culture containing all eight plasmids (3 biological replicates). (**D**) A comparison of the signals from individual RT-qPCR measurements of different strains and NGS analysis of a mixed culture reveals a linear trend (R^2^ = 0.88). The fold change between each construct and the weakest member of the set (P51 and pBBR1) is used to normalize data points to generate a trend. Fold changes are calculated pairwise, with either mean normalized copies of 16S rRNA (x-axis) or mean CLR transformed counts (y-axis), divided by the respective signal value of the weakest member. The P51 plasmids were removed to prevent data skewing as these presented much weaker expression.

To evaluate the barcoding signals arising from each construct, plasmids were individually transformed into *E. coli*. All eight cultures were pooled at equivalent titers to create a mixture containing all eight cat-RNA-v2, from which 3 aliquots were taken for downstream RNA extraction and analysis. For RT-qPCR measurements, barcoded-rRNA was quantified for each barcode and normalized to total 16S rRNA. The different promoters produced similar barcoded-rRNA signals across constructs with the two replication origins (Figure 6B), with the exception of the strongest promoter, which produced a ∼2-fold higher signal with the ColE1 origin. These results show that RAM can be multiplexed to report on promoters and origins of replication in parallel using one barcode per biological feature within *E. coli*.

To investigate whether NGS can be used to resolve eight biological features in *E. coli*, we sequenced the barcoded-rRNA from the samples that were evaluated using RT-qPCR. CLR-transformed sequencing data (Figure 6C) revealed a weak correlation with RT-qPCR signals (R^2^= 0.23) (Supplemental Figure 11). Based on our model predictions from mixed communities, we theorized that our limit of detection was too small to account for noise induced by inter-barcode variability with only a single orthogonal barcode. Therefore, the CLR-transformed counts of this mixed construct community were normalized by subtracting the centroid of the single promoter orthogonal barcode set as shown in Figure 4E. This normalization corrects for barcode-induced variability by simply adding or subtracting the known impacts from the results. Following this correction, the correlation (R^2^ = 0.88) between NGS and RT-qPCR signals improved substantially (Figure 6D). These results show that eight biological features, *i.e.*, 1 byte of information, can be resolved by NGS using eight orthogonal cat-RNA-v2 in *E. coli*, and they confirm that inter-barcode variability correction improves the quantitative comparisons using NGS-based RAM measurements.

### Storing 2.5 bytes in rRNA using five orthogonal cat-RNA-v2 per feature

To evaluate whether five orthogonal cat-RNA-v2 barcodes per transcriptional feature are sufficient to differentiate features from one another across multiple microbes, 40 constructs were created representing all combinations of the four promoters (P8, P34, P51, and P59) and two origins of replication (pBBR1 and ColE1), with a set of five orthogonal cat-RNA-v2 reporting on each promoter-origin combination. These constructs were transformed into *E. coli*, *V. natriegens*, and *P. putida* (Figure 7A). Based on the known host range of each origin, we expected RAM to resolve all eight features in *E. coli* and *V. natriegens*, but only four features in *P. putida*, since ColE1 is rarely functional in *Pseudomonas* (43). All 2.5 bytes of information were tested independently in RT-qPCR measurements and compared against RNA-sequencing results. Together, this experimental design was used to evaluate whether RAM can report on 20 biological features across three strains using NGS.

**Figure 7.**
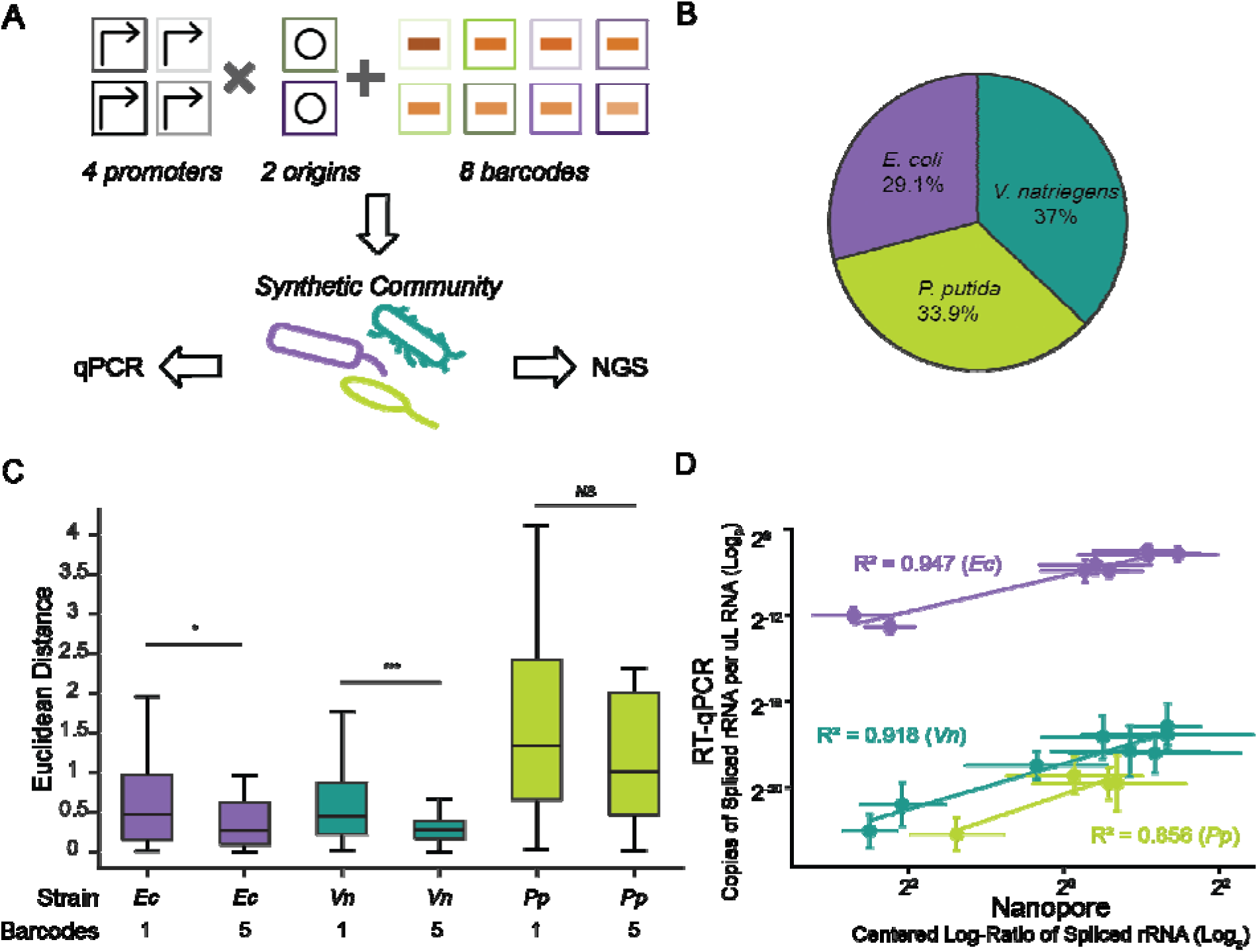
Recording 2.5 bytes of information across 16 rRNA from a trio of microbes. (**A**) The barcoded-rRNA signal was measured from *E. coli* (purple) and *V. natriegens* (teal) containing 40 plasmids (8 promoters, 2 origins, and 5 barcodes per promoter-origin combination) and *P. putida* (green) containing 20 plasmids (8 promoters, 1 origin, and 5 barcodes per combination). RT-qPCR was performed on each microbe containing the orthogonal cat-RNA-v2 used to report on each unique promoter-origin pair, while nanopore sequencing was used to report on extracted RNA from each strain that had been mixed (3 biological replicates). (**B**) The average abundances of barcoded-rRNA signals varied across organisms. (**C**) The Euclidean distances, derived from the pairwise distance between the CLR transformed counts of a given construct across replicates for each organism in the mixed community (n = 3 replicates, m = 28 unique pairwise distances). (**D**) A comparison of the RT-qPCR and the nanopore sequencing signals for plasmids representing each unique promoter-origin revealed linear trends across *E. coli* (R^2^ = 0.95), *V. natriegens* (R^2^ = 0.92), and *P. putida* (R^2^ = 0.86).

To assess the signals from the orthogonal plasmid pools (n = 5 barcodes per feature), we measured their signals in *E. coli* using RT-qPCR and compared them to prior measurements using a single cat-RNA-v2 barcode. A strong correlation (R^2^ 0.93) was observed between the orthogonal pooled barcodes and single barcode measurements, with no difference in rank order (Supplemental Figure 12). Orthogonal plasmid pools were next evaluated in *V. natriegens* and *P.* putida (Supplemental Figure 13A). All three species maintained a similar rank-order across the four promoters, with the exception of *P. putida*, in which the relative ranks of P51 and P08 (high and medium strength in *E. coli*, respectively) were swapped. These results establish a frame of reference for how barcoded-rRNA signals from pooled, orthogonal plasmids vary across the three species when reporting on the same biological features.

To analyze the barcoded-rRNA signals from a pooled mixture, we made two changes to the experimental design relative to prior measurements. First, RNA was extracted separately from each species and then mixed at equal ratios prior to the RT step to eliminate species bias. Second, we used long read nanopore sequencing to show that RAM is compatible with an additional sequencing platform. With this approach, all 100 unique barcode-rRNA pairs were detected (Supplemental Figure 13B), with similar counts observed across the three species (Figure 7B). To assess variability within this dataset, we evaluated the Euclidean distances across the five degenerate barcodes encoding each biological feature (Figure 7C). Compared to the prior community measurements, these distances were significantly lower across all three species (Wilcoxon sign-ranked t-test *E. coli* = 0.01; *V. natriegens* = 0.0005; *P. putida* = 0.07). These results show that using multiple orthogonal barcodes (n = 5) to report on each biological feature improves the consistency of barcoded-rRNA signals detected by NGS, consistent with our model predictions. Notably, this improvement did not reach statistical significance in *P. putida*, consistent with its higher baseline inter-barcode variability.

To investigate whether NGS can resolve 2.5 bytes of information using barcoded-rRNA across three microbes containing 100 unique barcoded-rRNA sequences, we compared NGS and RT-qPCR results directly. All three species showed strong correlations between NGS and RT-qPCR (R^2^: *E. coli* = 0.95; *V. natriegens* = 0.92; *P. putida* = 0.86) (Figure 7D). These results show that biological features representing 2.5 bytes of information can be monitored by sequencing barcoded-rRNA, supporting model predictions that estimated five orthogonal barcodes per feature are sufficient to resolve unique biological features within a synthetic community.

## DISCUSSION

We show that orthogonal cat-RNA barcodes can be designed to report on two plasmid-encoded features in parallel, promoter identity and origin of replication. Importantly, orthogonal cat-RNA-v2, which uses a tRNA-derived barcode, presents a more consistent signal compared to cat-RNA-v1, enabling the simultaneous detection of 1 byte of information in *E. coli* using RAM. We also show that 2.5 bytes of information can be read out from a community of three microbes transformed with a set of plasmids. We establish a quantitative model for predicting the number of orthogonal cat-RNA-v2 required to statistically resolve different plasmid-encoded features in parallel using NGS. While promoter activities can be read out in microbes using RNA sequencing (45), our approach offers an alternative route to obtain this information through targeted sequencing of short barcoded-rRNA amplicons, without requiring transcriptome-wide analysis. Our approach also allows more biological features to be monitored in parallel than fluorescent protein reporters (46), which have been used for parallel detection of promoter activities (47–49). At their best, these reporters can address only four biological features in parallel, due to spectral crosstalk (50). Thus, multiplexed RAM offers a high-throughput, one-pot approach for species-linked studies of plasmid-encoded genetic circuits in microbial communities.

Our measurements also extend the utility of RAM from studies of gene transfer to studies of transcriptional activity in communities (18, 19). While promoter activity can be inferred in communities using metatranscriptomics (51), taxonomic assignment of these signals depends on the availability of reference genomes and remains challenging in uncharacterized microbes (52). Our approach instead links promoter activity directly to host identity through the 16S rRNA itself, providing taxonomic information independent of reference genome availability. RAM could thus be used to probe sequence-function relationships in promoters across diverse microbes. Models have been developed for predicting promoter activities (40, 53, 54), but these models remain challenging to implement in high-throughput across microbial communities. By coupling different promoters to orthogonal cat-RNA-v2, RAM should be useful in future studies to evaluate the relative transcription rates of synthetic promoters across microbiomes, including unculturable microbes. Such studies could establish how sequence variation in sigma factors, which control transcription initiation, relates to the activity of synthetic promoters in different hosts (55), information that could in turn improve computational models of promoter activity (40, 53, 54).

Beyond promoters, multiplexed RAM should be useful for parallel, high-throughput studies of genetic parts in communities. For example, terminator strength could be assessed by placing synthetic terminators between a cat-RNA and its promoter, an approach similar to designs used with fluorescent protein reporters (56); the resulting abundance of barcoded-rRNA would then report on terminator strength across hosts in parallel. More broadly, any genetic part whose activity can be coupled to cat-RNA expression could be characterized using the same framework. Multiplexed RAM could also be extended to studies of mobile genetic element diversity within a single community, where libraries of mobile elements are each tagged with a unique orthogonal barcode, allowing the host range and relative mobility of many elements to be resolved in parallel using a single sequencing experiment. Multiplexed RAM may therefore serve as a foundational tool for comparing the activities of different genetic parts and mobile elements across hosts within communities, advancing microbiome engineering efforts (57, 58).

The variability observed with orthogonal cat-RNA highlights an important design consideration with multiplexed RAM. Our results show that orthogonal cat-RNA-v2 have more consistent signals than cat-RNA-v1 and should be used when performing multiplexed RAM. The larger variation observed with cat-RNA-v1 is thought to arise because its barcodes are more sensitive to RNA degradation than the tRNA-derived barcodes used in cat-RNA-v2 (59), which were designed to minimize RNAse sensitivity (31). Despite this improved consistency, cat-RNA-v2 still exhibited variability across species, with *P. putida* showing the highest variability. The underlying cause of this species-to-species variability is not known. One possibility is that the tRNA-derived barcodes undergo post-transcriptional modifications that alter their stabilities and vary across microbes (60, 61). Another possibility is that the sequence variation within the barcodes differentially affects binding to proteins involved in post-transcriptional modification or aminoacyl tRNA synthetases (62, 63). Future studies could probe these possibilities by evaluating the barcoding of hundreds of orthogonal cat-RNA in parallel across many different microbes to identify sequence features that correlate with high variability. Such experiments can be performed in microbes with and without specific tRNA-modifying proteins and RNAses to establish their roles in regulating barcode stability.

## Data Availability

The data underlying this article are available in the article and in its online supplementary material.

## Supplementary Data statement

Supplementary Data are available at *NAR* Online.

## Author Contributions Statement

Conceptualization: M.J.D, L.F., L.B.S., and J.J.S; Data Curation: M.J.D and L.F.; Formal Analysis: M.J.D, L.F., J.C., L.B.S., and J.J.S; Funding Acquisition: J.C., L.B.S., and J.J.S; Investigation: M.J.D, L.F., J.C., L.B.S., and J.J.S; Methodology: M.J.D, L.F., L.K., J.C., L.B.S., and J.J.S; Project Administration: L.B.S. and J.J.S; Supervision: L.B.S. and J.J.S; Validation: M.J.D and L.F.; Visualization: M.J.D, L.F., J.C., and L.B.S.; Writing - Original draft: M.J.D, L.F., L.B.S., and J.J.S; and Writing – review & editing: M.J.D, L.F., L.K., J.C., L.B.S., and J.J.S

## Funding

This work was supported by the following funding agencies: National Science Foundation (grant no. 2227526 to J.J.S. and J.C.; CAREER grant no. 2237052 to L.B.S.; CAREER grant no. 2237512 to J.C.) and the Army Research Office (W911NF-24-2-0073 to J.J.S., J.C. and L.B.S.). The views and conclusions contained in this document are those of the authors and should not be interpreted as representing the official policies, either expressed or implied, of the Army Research Office or the US Government. The US Government is authorized to reproduce and distribute reprints for Government purposes, notwithstanding any copyright notation.

## Conflict of interest

The authors have no conflict of interest to declare.

## Notes

### Competing Interest Statement

The authors have declared no competing interest.

## REFERENCES

1. Gilliot, P.-A. and Gorochowski, T.E. (2020) Sequencing enabling design and learning in synthetic biology. Current Opinion in Chemical Biology, 58, 54–62.

2. Wu, X., Yang, X., Dai, Y., Zhao, Z., Zhu, J., Guo, H. and Yang, R. (2024) Single-cell sequencing to multi-omics: technologies and applications. Biomark Res, 12, 110.

3. Mee, M.T. and Wang, H.H. (2012) Engineering ecosystems and synthetic ecologies. Molecular BioSystems, 8, 2470–2483.

4. Wong, A.S.L., Choi, G.C.G., Cui, C.H., Pregernig, G., Milani, P., Adam, M., Perli, S.D., Kazer, S.W., Gaillard, A., Hermann, M., et al. (2016) Multiplexed barcoded CRISPR-Cas9 screening enabled by CombiGEM. Proc. Natl. Acad. Sci. U.S.A., 113, 2544–2549.

5. Guo, X., Chavez, A., Tung, A., Chan, Y., Kaas, C., Yin, Y., Cecchi, R., Garnier, S.L., Kelsic, E.D., Schubert, M., et al. (2018) High-throughput creation and functional profiling of DNA sequence variant libraries using CRISPR–Cas9 in yeast. Nat Biotechnol, 36, 540–546.

6. Larson, M.H., Gilbert, L.A., Wang, X., Lim, W.A., Weissman, J.S. and Qi, L.S. (2013) CRISPR interference (CRISPRi) for sequence-specific control of gene expression. Nat Protoc, 8, 2180–2196.

7. Liu, Y., Wan, X. and Wang, B. (2019) Engineered CRISPRa enables programmable eukaryote-like gene activation in bacteria. Nat Commun, 10, 3693.

8. Zik, J.J., Price, M.N., Mayra, K.H.A., Santoso, A.A., Arkin, A.P., Deutschbauer, A.M. and Sham, L.-T. (2025) Dual transposon sequencing profiles the genetic interaction landscape in bacteria. Science, 389, eadt7685.

9. Loveless, T.B., Carlson, C.K., Dentzel Helmy, C.A., Hu, V.J., Ross, S.K., Demelo, M.C., Murtaza, A., Liang, G., Ficht, M., Singhai, A., et al. (2025) Open-ended molecular recording of sequential cellular events into DNA. Nat Chem Biol, 21, 512–521.

10. Kalhor, R., Kalhor, K., Mejia, L., Leeper, K., Graveline, A., Mali, P. and Church, G.M. (2018) Developmental barcoding of whole mouse via homing CRISPR. Science, 361, eaat9804.

11. Rai, K., O’Connell, R.W., Piepergerdes, T.C., Wang, Y., Brown, L.B.C., Samra, K.D., Wilson, J.A., Lin, S., Zhang, T.H., Ramos, E.M., et al. (2026) Ultra-high-throughput mapping of genetic design space. Nature, 650, 1035–1044.

12. Rubin, B.E., Diamond, S., Cress, B.F., Crits-Christoph, A., Lou, Y.C., Borges, A.L., Shivram, H., He, C., Xu, M., Zhou, Z., et al. (2021) Species- and site-specific genome editing in complex bacterial communities. Nat Microbiol, 7, 34–47.

13. Saliba, A.-E., Westermann, A.J., Gorski, S.A. and Vogel, J. (2014) Single-cell RNA-seq: advances and future challenges. Nucleic Acids Res, 42, 8845–8860.

14. Huang, D., Ma, N., Li, X., Gou, Y., Duan, Y., Liu, B., Xia, J., Zhao, X., Wang, X., Li, Q., et al. (2023) Advances in single-cell RNA sequencing and its applications in cancer research. J Hematol Oncol, 16, 98.

15. Zilionis, R., Nainys, J., Veres, A., Savova, V., Zemmour, D., Klein, A.M. and Mazutis, L. (2017) Single-cell barcoding and sequencing using droplet microfluidics. Nat Protoc, 12, 44–73.

16. Klein, A.M., Mazutis, L., Akartuna, I., Tallapragada, N., Veres, A., Li, V., Peshkin, L., Weitz, D.A. and Kirschner, M.W. (2015) Droplet Barcoding for Single-Cell Transcriptomics Applied to Embryonic Stem Cells. Cell, 161, 1187–1201.

17. Cetnar, D.P., Hossain, A., Vezeau, G.E. and Salis, H.M. (2024) Predicting synthetic mRNA stability using massively parallel kinetic measurements, biophysical modeling, and machine learning. Nat Commun, 15, 9601.

18. Kalvapalle, P.B., Staubus, A., Dysart, M.J., Gambill, L., Reyes Gamas, K., Lu, L.C., Silberg, J.J., Stadler, L.B. and Chappell, J. (2026) Information storage across a microbial community using universal RNA barcoding. Nat Biotechnol, 44, 269–276.

19. LaTurner, Z.W., Dysart, M.J., Schwartz, S.K., Zeng, E., Chappell, J., Silberg, J.J. and Stadler, L.B. (2026) Cross-order detection of bacteriophage transduction in microbial communities using RNA barcoding. Nat Commun, 17, 4308.

20. Reyes Gamas, K., Seamons, T.R., Dysart, M.J., Fang, L., Chappell, J., Stadler, L.B. and Silberg, J.J. (2025) Controlling the Taxonomic Composition of Biological Information Storage in 16S rRNA. ACS Synth. Biol., 14, 3530–3542.

21. Wei, Y. and Lin, F. (2025) Barcodes based on nucleic acid sequences: Applications and challenges (Review). Mol Med Rep, 32, 1–21.

22. Sachs, A.B. (1993) Messenger RNA degradation in eukaryotes. Cell, 74, 413–421.

23. Boo, S.H. and Kim, Y.K. (2020) The emerging role of RNA modifications in the regulation of mRNA stability. Exp Mol Med, 52, 400–408.

24. Chen, F., Cocaign-Bousquet, M., Girbal, L. and Nouaille, S. (2022) 5’UTR sequences influence protein levels in Escherichia coli by regulating translation initiation and mRNA stability. Front. Microbiol., 13, 1088941.

25. Salis, H.M., Mirsky, E.A. and Voigt, C.A. (2009) Automated design of synthetic ribosome binding sites to control protein expression. Nat Biotechnol, 27, 946–950.

26. Anderson, P., Monforte, J., Tritz, R., Nesbitt, S., Hearst, J. and Hamper, A. (1994) Mutagenesis of the hairpin ribozyme. Nucl Acids Res, 22, 1096–1100.

27. Young, B., Herschlag, D. and Cech, T.R. (1991) Mutations in a nonconserved sequence of the Tetrahymena ribozyme increase activity and specificity. Cell, 67, 1007–1019.

28. Cetnar, D.P. and Salis, H.M. (2021) Systematic Quantification of Sequence and Structural Determinants Controlling mRNA stability in Bacterial Operons. ACS Synth. Biol., 10, 318–332.

29. Deutscher, M.P. (2006) Degradation of RNA in bacteria: comparison of mRNA and stable RNA. Nucleic Acids Research, 34, 659–666.

30. Jackowiak, P., Nowacka, M., Strozycki, P.M. and Figlerowicz, M. (2011) RNA degradome--its biogenesis and functions. Nucleic Acids Research, 39, 7361–7370.

31. Karinje, L., Silberg, J.J., Chappell, J. and Stadler, L.B. (2026) Rational engineering enhances the signal and modularity of an RNA barcoding technology to track gene transfer in microbiomes. 10.64898/2026.04.20.719664.

32. Lorenz, R., Bernhart, S.H., Höner Zu Siederdissen, C., Tafer, H., Flamm, C., Stadler, P.F. and Hofacker, I.L. (2011) ViennaRNA Package 2.0. Algorithms Mol Biol, 6, 26.

33. Gibson, D.G., Young, L., Chuang, R.-Y., Venter, J.C., Hutchison, C.A. and Smith, H.O. (2009) Enzymatic assembly of DNA molecules up to several hundred kilobases. Nat Methods, 6, 343–345.

34. Engler, C., Kandzia, R. and Marillonnet, S. (2008) A One Pot, One Step, Precision Cloning Method with High Throughput Capability. PLoS ONE, 3, e3647.

35. Fernandes, A.D., Macklaim, J.M., Linn, T.G., Reid, G. and Gloor, G.B. (2013) ANOVA-Like Differential Expression (ALDEx) Analysis for Mixed Population RNA-Seq. PLoS ONE, 8, e67019.

36. Aitchison, J. (1994) Principles of compositional data analysis. In Institute of Mathematical Statistics Lecture Notes - Monograph Series. Institute of Mathematical Statistics, Hayward, CA, pp. 73–81.

37. Bolyen, E., Rideout, J.R., Dillon, M.R., Bokulich, N.A., Abnet, C.C., Al-Ghalith, G.A., Alexander, H., Alm, E.J., Arumugam, M., Asnicar, F., et al. (2019) Reproducible, interactive, scalable and extensible microbiome data science using QIIME 2. Nat Biotechnol, 37, 852–857.

38. Che, A.J. and Knight, T.F. (2010) Engineering a family of synthetic splicing ribozymes. Nucleic Acids Research, 38, 2748–2755.

39. Estrada, K., Garciarrubio, A. and Merino, E. (2024) Unraveling the plasticity of translation initiation in prokaryotes: Beyond the invariant Shine-Dalgarno sequence. PLoS ONE, 19, e0289914.

40. LaFleur, T.L., Hossain, A. and Salis, H.M. (2022) Automated model-predictive design of synthetic promoters to control transcriptional profiles in bacteria. Nat Commun, 13, 5159.

41. Jahn, M., Vorpahl, C., Hübschmann, T., Harms, H. and Müller, S. (2016) Copy number variability of expression plasmids determined by cell sorting and Droplet Digital PCR. Microb Cell Fact, 15, 211.

42. Kovach, M.E., Elzer, P.H., Steven Hill, D., Robertson, G.T., Farris, M.A., Roop, R.M. and Peterson, K.M. (1995) Four new derivatives of the broad-host-range cloning vector pBBR1MCS, carrying different antibiotic-resistance cassettes. Gene, 166, 175–176.

43. Ares-Arroyo, M., Rocha, E.P.C. and Gonzalez-Zorn, B. (2021) Evolution of ColE1-like plasmids across γ-Proteobacteria: From bacteriocin production to antimicrobial resistance. PLoS Genet, 17, e1009919.

44. Schmidt, L. and Inselburg, J. (1982) ColE1 copy number mutants. J Bacteriol, 151, 845–854.

45. Bullard, J.H., Purdom, E., Hansen, K.D. and Dudoit, S. (2010) Evaluation of statistical methods for normalization and differential expression in mRNA-Seq experiments. BMC Bioinformatics, 11, 94.

46. Han, J., Xia, A., Huang, Y., Ni, L., Chen, W., Jin, Z., Yang, S. and Jin, F. (2019) Simultaneous Visualization of Multiple Gene Expression in Single Cells Using an Engineered Multicolor Reporter Toolbox and Approach of Spectral Crosstalk Correction. ACS Synth. Biol., 8, 2536–2546.

47. Cox, R.S., Dunlop, M.J. and Elowitz, M.B. (2010) A synthetic three-color scaffold for monitoring genetic regulation and noise. J Biol Eng, 4, 10.

48. Parrello, D., Mustin, C., Brie, D., Miron, S. and Billard, P. (2015) Multicolor Whole-Cell Bacterial Sensing Using a Synchronous Fluorescence Spectroscopy-Based Approach. PLoS ONE, 10, e0122848.

49. Wu, F., Van Rijn, E., Van Schie, B.G.C., Keymer, J.E. and Dekker, C. (2015) Multi-color imaging of the bacterial nucleoid and division proteins with blue, orange, and near-infrared fluorescent proteins. Front. Microbiol., 6.

50. Dickinson, M.E., Bearman, G., Tille, S., Lansford, R. and Fraser, S.E. (2001) Multi-Spectral Imaging and Linear Unmixing Add a Whole New Dimension to Laser Scanning Fluorescence Microscopy. BioTechniques, 31, 1272–1278.

51. Zhang, Y., Thompson, K.N., Branck, T., Yan Yan, Nguyen, L.H., Franzosa, E.A. and Huttenhower, C. (2021) Metatranscriptomics for the Human Microbiome and Microbial Community Functional Profiling. Annu. Rev. Biomed. Data Sci., 4, 279–311.

52. Bharti, R. and Grimm, D.G. (2021) Current challenges and best-practice protocols for microbiome analysis. Briefings in Bioinformatics, 22, 178–193.

53. Cassiano, M.H.A. and Silva-Rocha, R. (2020) Benchmarking Bacterial Promoter Prediction Tools: Potentialities and Limitations. mSystems, 5, e00439–20.

54. Huang, Y.-K., Yu, C.-H. and Ng, I.-S. (2024) Precise strength prediction of endogenous promoters from Escherichia coli and J-series promoters by artificial intelligence. Journal of the Taiwan Institute of Chemical Engineers, 160, 105211.

55. Feklístov, A., Sharon, B.D., Darst, S.A. and Gross, C.A. (2014) Bacterial Sigma Factors: A Historical, Structural, and Genomic Perspective. Annu. Rev. Microbiol., 68, 357–376.

56. Chen, Y.-J., Liu, P., Nielsen, A.A.K., Brophy, J.A.N., Clancy, K., Peterson, T. and Voigt, C.A. (2013) Characterization of 582 natural and synthetic terminators and quantification of their design constraints. Nat Methods, 10, 659–664.

57. Diao, J., Tian, Y., Park, S., Cho, S.-M. and Moon, T.S. (2025) Engineering microbial consortia for mixed plastic upcycling. Nat Commun, 17, 637.

58. USDOE Office of Science (SC), Biological and Environmental Research (BER), US Department of Energy (USDOE), Washington, DC (United States). Office of Science, Cregger, M., Silberg, J., Bonito, G., Henry, C., Hofmockel, K., O’Malley, M. and You, L. (2025) Engineering Microbial Communities.

59. Tejada-Arranz, A., De Crécy-Lagard, V. and De Reuse, H. (2020) Bacterial RNA Degradosomes: Molecular Machines under Tight Control. Trends in Biochemical Sciences, 45, 42–57.

60. De Crécy-Lagard, V. and Jaroch, M. (2021) Functions of Bacterial tRNA Modifications: From Ubiquity to Diversity. Trends in Microbiology, 29, 41–53.

61. Lorenz, C., Lünse, C. and Mörl, M. (2017) tRNA Modifications: Impact on Structure and Thermal Adaptation. Biomolecules, 7, 35.

62. Björk, G.R. and Hagervall, T.G. (2014) Transfer RNA Modification: Presence, Synthesis, and Function. EcoSal Plus, 6, 10.1128/ecosalplus.ESP-0007–2013.

63. Giegé, R. and Springer, M. (2016) Aminoacyl-tRNA Synthetases in the Bacterial World. EcoSal Plus, 7, 10.1128/ecosalplus.ESP-0002–2016.

