## Supplemental Figures 1-13; Supplemental Tables 1-6 for "Storing >1 byte of information in 16S ribosomal RNA using orthogonal trans-splicing ribozymes"

### TABLE OF CONTENTS


| Item | Title | Page |
|---|---|---|
| Figure S1 | Using Euclidean distances to analyze cat-RNA NGS data | 3 |
| Figure S2 | Strategy to generate a library of orthogonal catRNA-v1 | 4 |
| Figure S3 | Data analysis pipeline for NGS | 5 |
| Figure S4 | Signals from cat-RNA-v1 transcribed from 23 promoters | 6 |
| Figure S5 | Cat-RNA-v1 signals vary across P1 and P18 promoters | 7 |
| Figure S6 | Abundances of catalytically-dead cat-RNA-v1 | 8 |
| Figure S7 | Rank order of barcoded-rRNA and inactive cat-RNA signals | 9 |
| Figure S8 | Species abundance in the synthetic community | 10 |
| Figure S9 | Rank order of barcoded-rRNA signals across microbes | 11 |
| Figure S10 | Effect of mixing species on signal variability in each microbe | 12 |
| Figure S11 | Raw signals for 1 byte measurement in *E. coli* | 13 |
| Figure S12 | Pooled and single barcode RT-qPCR comparison | 14 |
| Figure S13 | RT-qPCR and NGS from 8 features across 3 microbes | 15 |
| Table S1 | List of plasmids | 16 |
| Table S2 | Promoters used to transcribe cat-RNA | 17 |
| Table S3 | Sequence variation generated within cat-RNA barcodes | 18 |
| Table S4 | The 16S rRNA sequences in barcoded-rRNA amplicons | 18 |
| Table S5 | Primers used for RT-qPCR and NGS | 19 |
| Table S6 | Viennafold predictions | 21 |

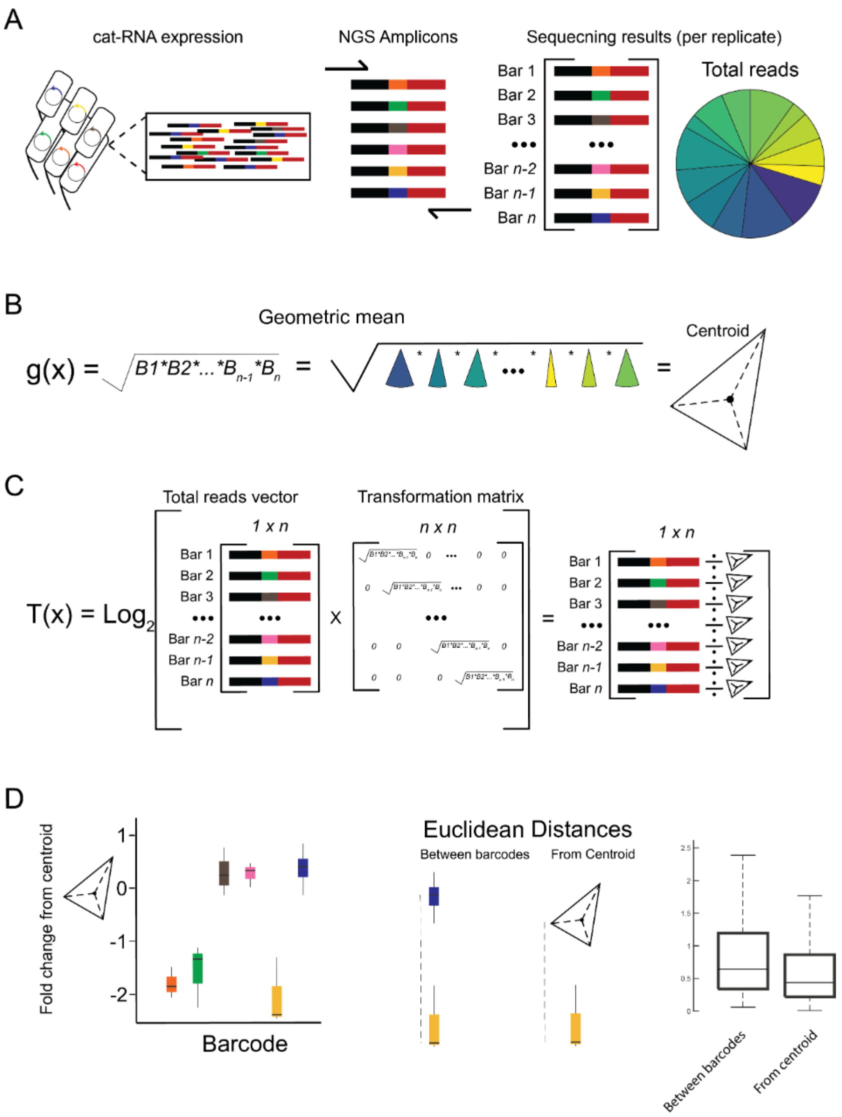


**Supplemental Figure 1. Using Euclidean distances to analyze cat-RNA NGS data.** (**A**) A cat-RNA experiment is read out using NGS, creating A-vector space consisting of the total number of reads per barcode. The vector space is the equivalent of the frequency of reads, which is a simplex dataspace. (**B**) To normalize the dataset and transform it into the Euclidean domain, each element of the dataset is divided by the geometric mean. This value is the central tendency of the simplex, which anchors the relative proportions of reads into the Euclidean space. (**C**) A logarithm is taken of these values to observe the differences as fold change distances from the central tendency. (**D**) For cat-RNA datasets, the transformed dataset exists on a single plane and therefore the Euclidean distance, the dimension specific distance one point is from another, simplifies to the magnitude of the difference between the two centered log-ratio transformed points. There are two unique differences, which are preserved in the transformation. The distance from the centroid, and the pairwise distance to another element of the transformed simplex.

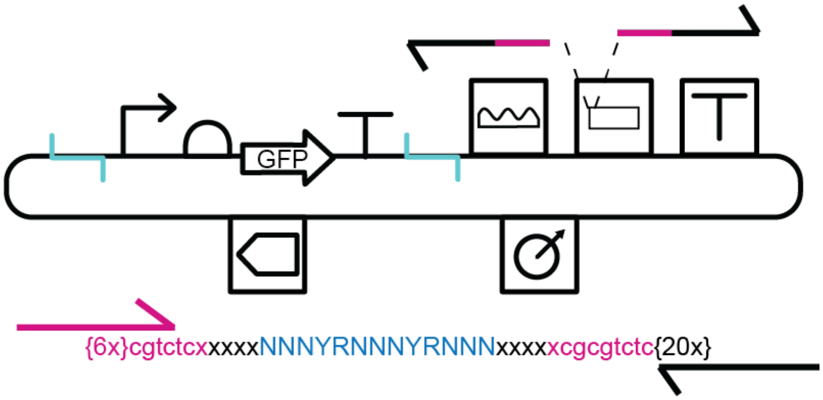


**Supplemental Figure 2. Strategy to generate a library of orthogonal catRNA-v1.** Example of template used to generate random barcode sequences for cat-RNA-v1. An oligo (bottom) with thirteen random oligos of a specific pattern (blue), was flanked by restriction enzyme sites (pink). When extended to produce a dsDNA fragment, the barcode sequences were inserted into a linearized cat-RNA-v1 vector (top) in between the probe binding site and the RT-primer binding.

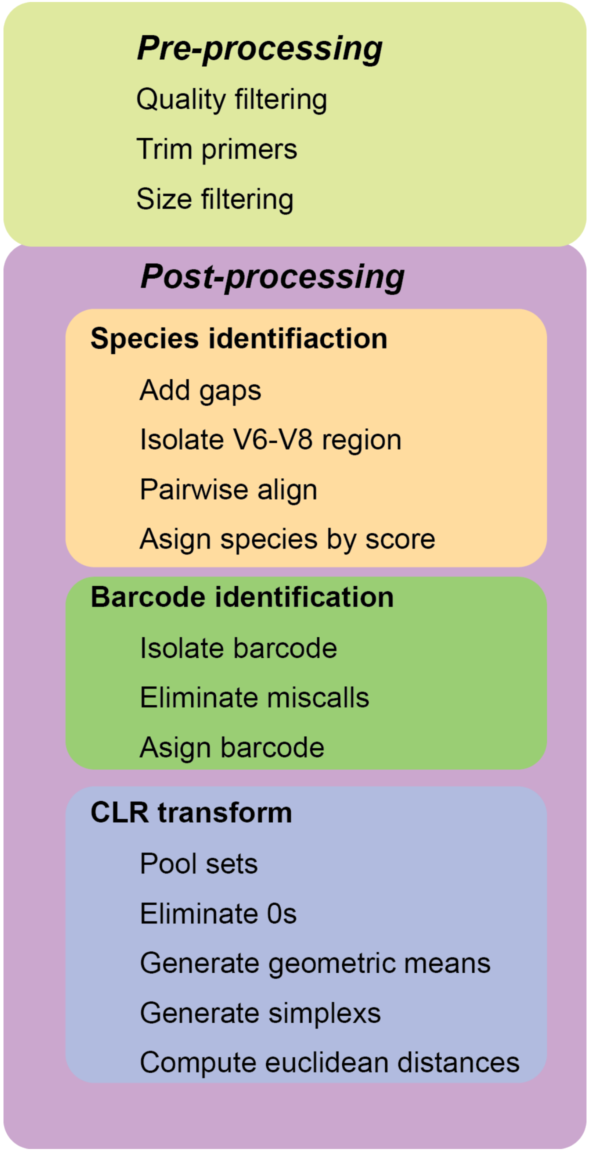


**Supplemental Figure 3. Data analysis pipeline for NGS**. Workflow showing data processing to quantify barcoding. Built in function FASTQread() was used to isolate sequence and quality scores for pre-processing. Custom script analysis was split into three function types: Species identification, Barcode identification, and CLR transformation. Statics were computed only on transformed datasets. Code available on Github.

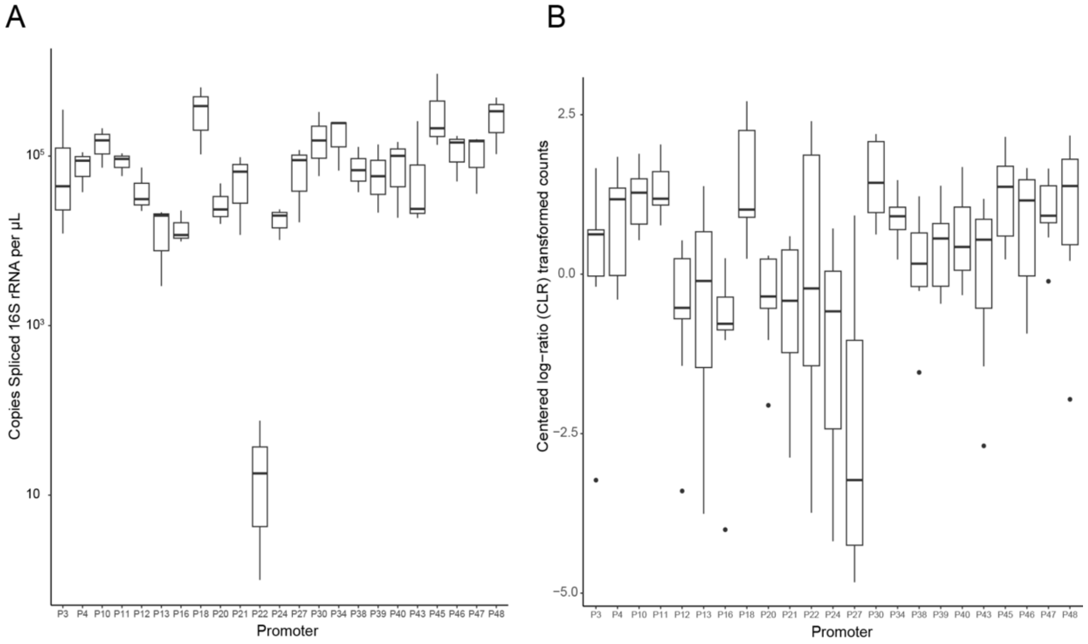


**Supplemental Figure 4. Signals from cat-RNA-v1 transcribed from 23 promoters.** Promoters are given numbers and represented by an uppercase P. Promoters that did not have RT-qPCR or NGS data for all three barcodes were excluded from these results. **(A)** RT-qPCR results on copies of barcoded-rRNA per µL of RNA extract. Orthogonal barcodes expressed by the same promoter were treated as if they were biological replicates (*n*= 3). **(B)** NGS results from a pooled community of cat-RNA-v1 transcribed by trios of orthogonal cat-RNA (*n*=3). All simplex datasets from NGS were transformed into the Euclidean space by centered log-ratio (CLR) transformation for statistical analysis. To adjust for mixing errors, barcoded-rRNA CLR counts were normalized by their paired DNA plasmid CLR counts, to correct for over or underrepresented of a given barcode within the total community. Orthogonal barcodes expressed by the same promoter were treated as biological replicates (*n*=3). Resulting boxplots are composed of 9 CLR transformed counts, comprising 3 orthogonal barcodes from 3 sequencing runs.

A

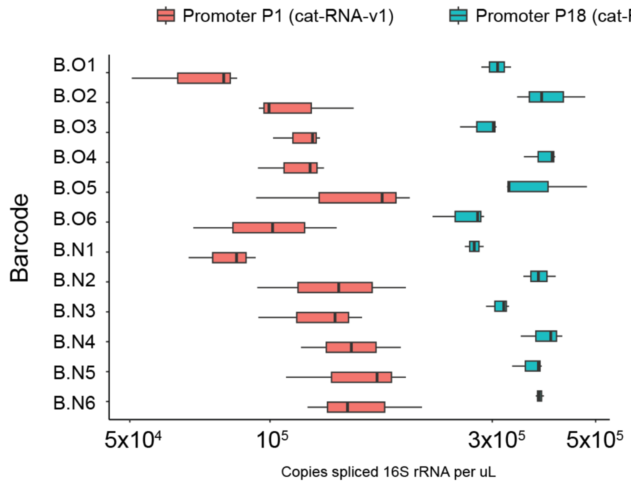


B

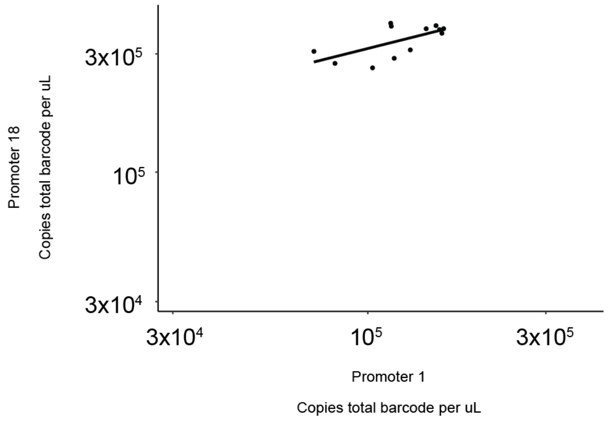


**Supplemental Figure 5. Cat-RNA-v1 signals vary across P1 and P18 promoters.** Primer probe set targets a region of the barcode that exists in both barcoded-rRNA and unspliced ribozyme transcripts. **(A)** cat-RNA-v1 RT-qPCR results on twelve unique barcodes (n=3). Each barcode was expressed by both promoter 1 (yellow) and promoter 18 (cyan). CT values are converted into copies per uL based on a standard curve for total barcode. **(B)** The correlation between the copies of total barcode transcribed by either promoter 1 or promoter 18, with a linear regression of 0.27

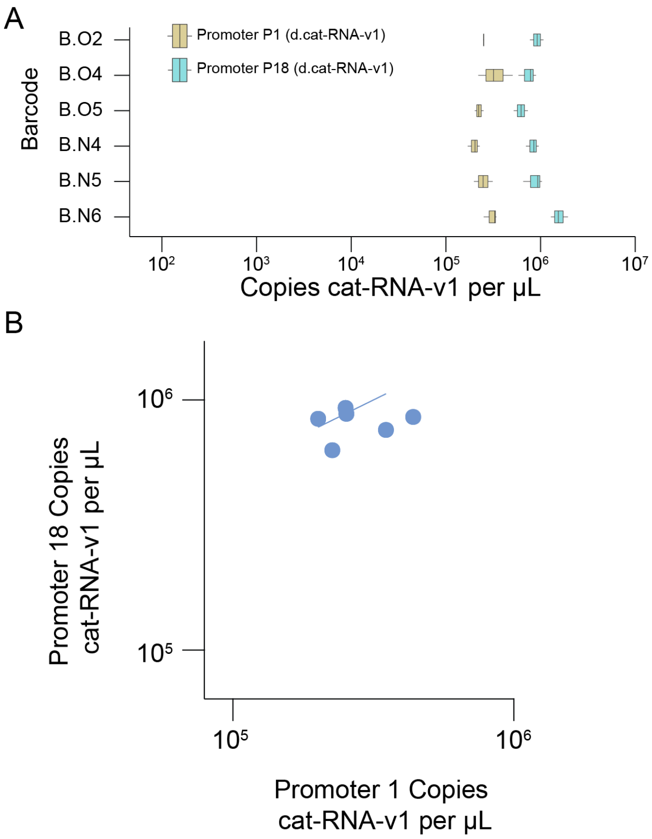


**Supplemental Figure 6. Abundances of catalytically-dead cat-RNA-v1.** (**A**) The d.cat-RNA-v1 RT-qPCR results of six unique barcodes (n=3). Each barcode was expressed by both promoter 1 (yellow) and promoter 18 (cyan). CT values are converted into copies per uL based on a standard curve. (**B**) The correlation between the copies of d.cat-RNA-v1 transcribed by either promoter 1 or promoter 18, with linear regression yielding an $R^2$ of 0.37

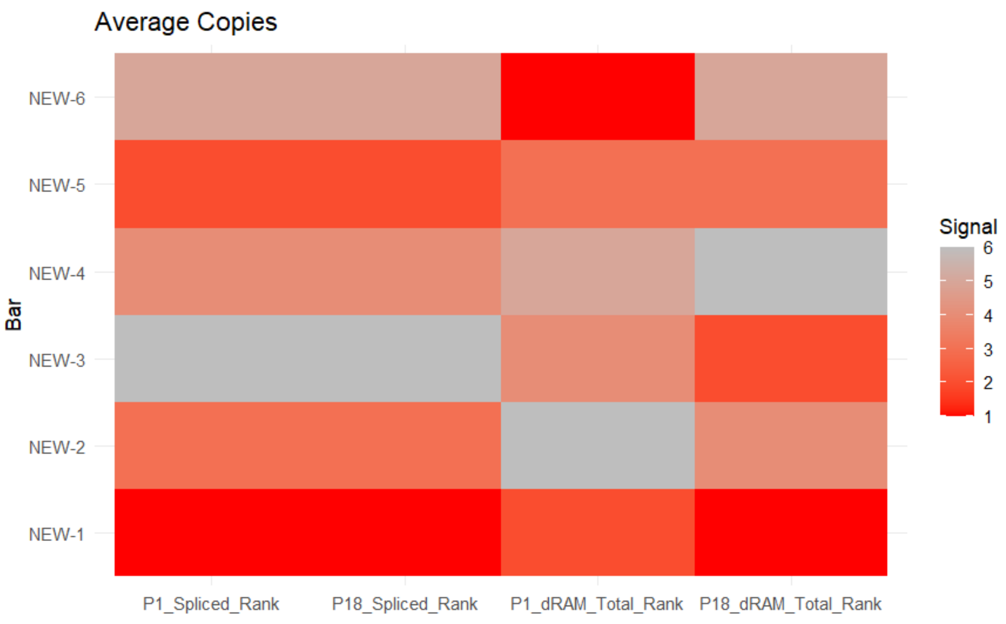


**Supplemental Figure 7. Effect of cat-RNA inactivation on rank order of barcoded-rRNA.** The rank order of the barcoded-rRNA abundances arising from six cat-RNA-v1 transcribed using P1 (P1_Sliced_Rank) and P18 (P18_Sliced_Rank) is compared with the cat-RNA-v1 abundances for inactive variants (dRAM) expressed using the same promoters. Signal represents the rank order strength of RT-qPCR results as absolute average copies observed.

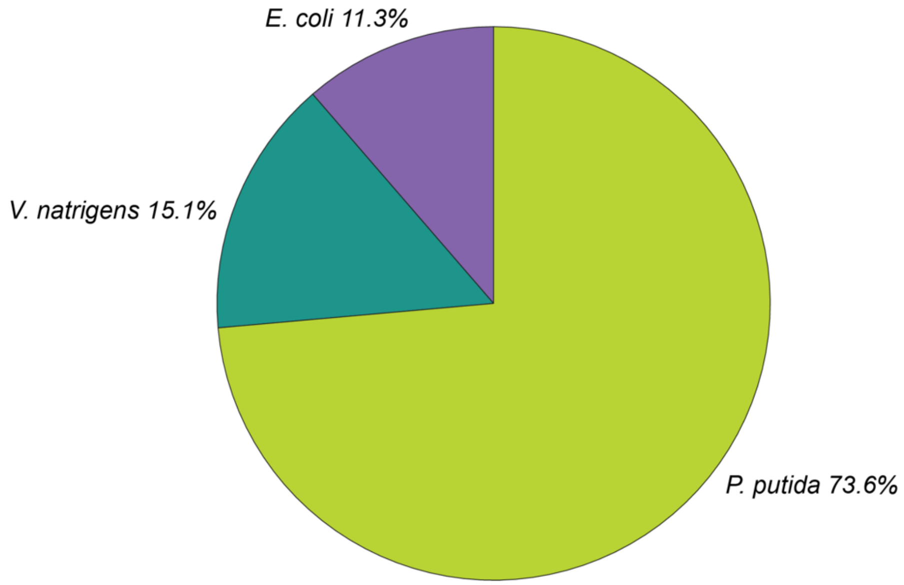


**Supplemental Figure 8. Species abundance in the synthetic community.** Genomic DNA was extracted from a frozen pellet of the constructed synthetic community as described in the methods. Amplicons of the 16S rRNA gene were amplified using NGS primers as listed in Supplemental Table 2 and with conditions as described in the methods. Results are the frequency of total raw counts after each read was merged and pairwise compared to the three reference 16S rRNA gene sequences.

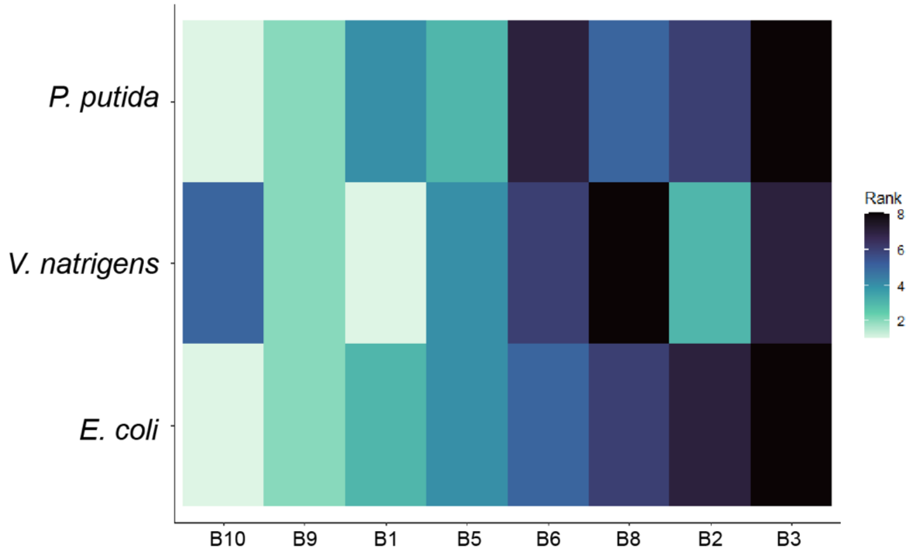


**Supplemental Figure 9. Rank order of barcoded-rRNA signals across microbes.** Rank order of signal strength as determined by NGS analysis of cat-RNA-v2 barcoded reads from 8 orthogonal barcodes expressed in a synthetic community by a single construct. X-axis represents barcodes, from least observed to most observed, based on observed trends in *E. coli*. Y- axis delineates species.

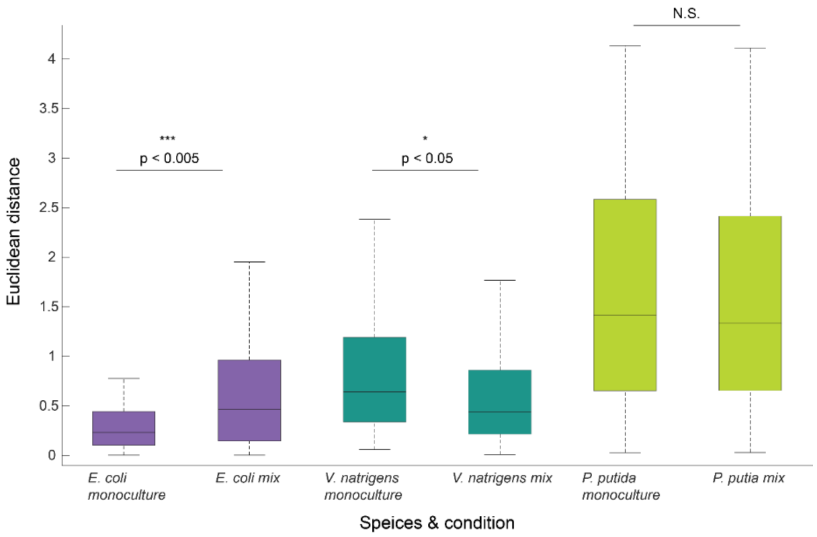


**Supplemental Figure 10. Effect of mixing species on signal variability in each microbe.** Monocultures are composed of each species pooled by itself and mixed represent species specific reads from the synthetic community. Boxplots are composed of the Euclidean distances (n = 225 *E. coli pure*; n = 84 all other sets). Non-parametric Wilcoxon sign-ranked t-test was used to compare each monoculture dataset with the respective species data from the synthetic community.

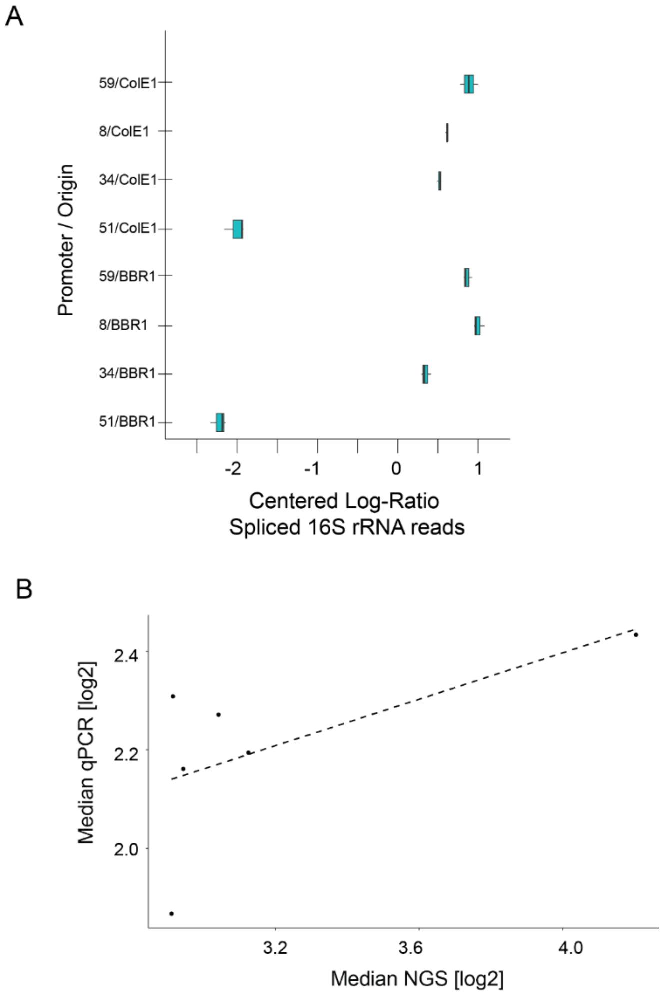


**Supplemental Figure 11. Raw signals for 1 byte measurement in *E. coli*.** (**A**) Using the same x-axis as the RT-qPCR plot from figure 3B, the unadjusted NGS results were transformed by the centered log-ratio of a given sequencing dataset for replicate normalization. These values are reported as Centered log-ratio barcoded-rRNA, which is a representation of the frequency of a given construct with respect to the geometric mean (n=3) (**B**) A correlation plot observing the differences between RT-qPCR and unadjusted NGS data. The fold change between each construct and the weakest member of the set (P51 - pBBR1) is used to evaluate the trend. Fold changes are calculated arithmetically. Both P51 constructs were removed from the correlation to prevent skewed data by the significantly weaker expression seen in these two constructs, and also being the basis of comparison. The linear regression reported an $R^2$ value of 0.23 suggesting a weak correlation between RT-qPCR and NGS results.

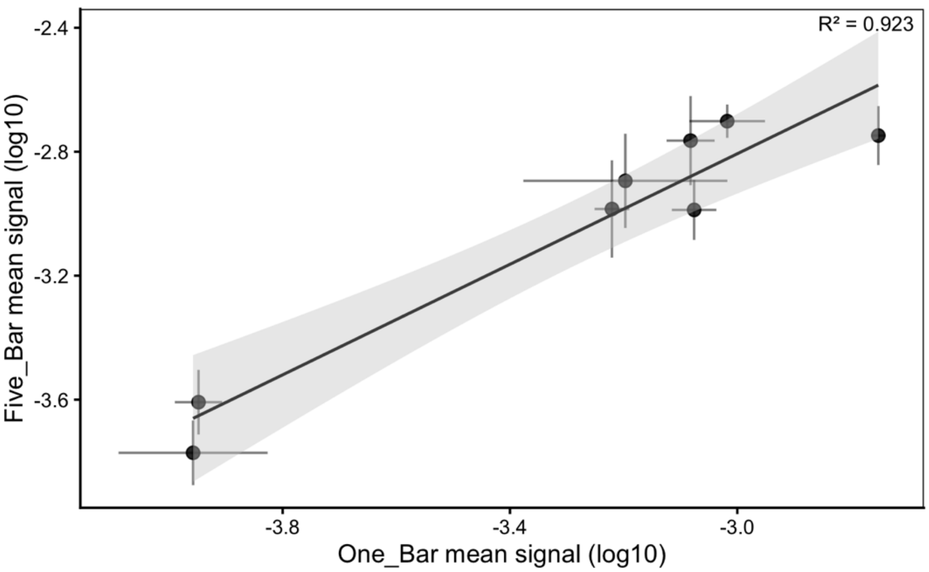


**Supplemental Figure 12. Pooled and single barcode RT-qPCR comparison.** Individual barcodes on the 8 unique constructs were compared against the set of five degenerate barcodes assigned to each construct. Values, as points, represented the log10 transformed ratio of copies of native 16S rRNA to spliced 16S rRNA for a sample as determined by RT-qPCR for each construct (n = 3 replicates). X-axis represents a single barcode and the Y-axis represents the set of five barcodes per construct. $R^2$ was determined by linear regression fit.

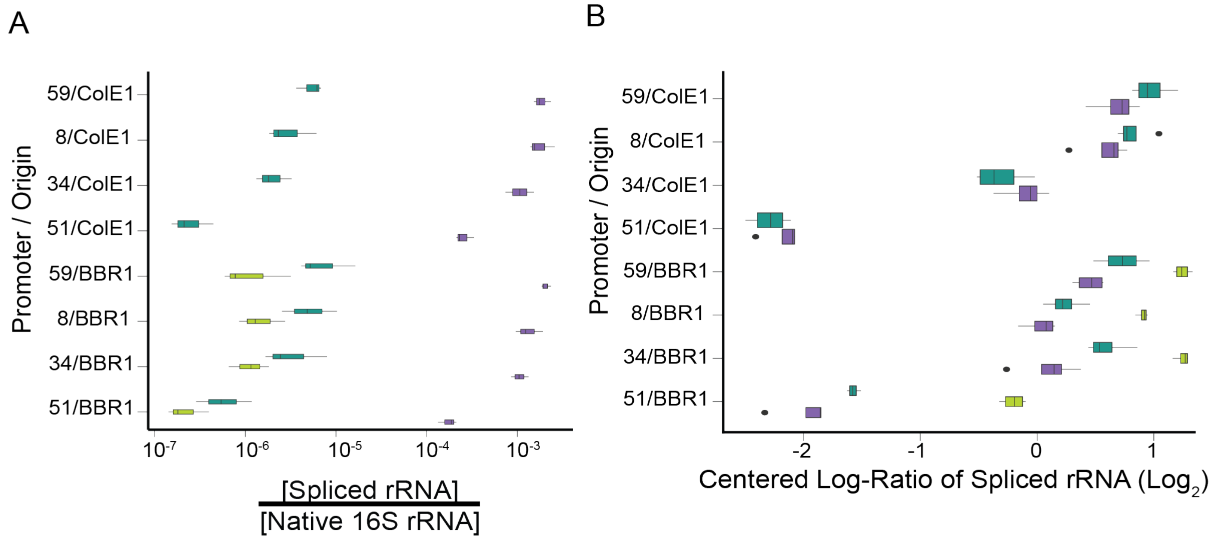


**Supplemental Figure 13. RT-qPCR and NGS from 8 features across 3 microbes.** (**A**) RT-qPCR analysis of barcoded-rRNA signals from each set of orthogonal cat-RNA. The barcoded-rRNA signal for each sample was normalized to total 16S rRNA. Three biological replicates were performed for each measurement. (**B**) For each unique construct, the relative abundance of sequence counts is shown using centered log-ratios (n=3). Barcodes are ordered in increasing promoter strength as determined in the previous experiment, separated into two groups by the construct backbone.

**Supplemental Table 1. List of plasmids.** For each, we note the use, the relevant data, and the architecture and features, including: (i) selectable marker, (ii) origin of replication, (iii) promoter, (iv) translation initiation sequences, (v) open reading frames, and (vi) ribozyme details.

| Category | Name | Name | Use | Figure | Architecture |
|---|---|---|---|---|---|
| *Barcode stability + Orthogonal barcode signal variation tests* | pPK076 | cat-RNA-v1 template | Basis of all cat-RNA-v1 orthogonal plasmids | *Not Shown* | PcymR - cat-RNA-v1-tVoigtS4 - KanR.II - pBBR1 |
| | pMJD089 | Library of cat-RNA-v1 | Library of synthetic promoters expressing trios of cat-RNA-v1 orthogonal barcodes | 2G,S4 | [Synthetic promoter]-cat-RNA-v1-tVoigtS4 - KanR.II - pBBR1 |
| | pSPL001-pSPL012 | 12 barcodes P1 | Average signal cat-RNA-v1 | 2C,2E,S5 | P1- cat-RNA-v1-tVoigtS4 - KanR.II - pBBR1 |
| | pSPL013-pSP024 | 12 barcodes P18 | Strong signal cat-RNA-v1 | 2C,2E,S5 | P18 - cat-RNA-v1-tVoigtS4 - KanR.II - pBBR1 |
| | pSPL025 | d.RAM cat-RNA-v1 template | Template plasmid for deactivated cat-RNA-v1 | S6 | PcymR - d.cat-RNA-v1-tVoigtS4 - KanR.II - pBBR1 |
| | pLNK007 | cat-RNA-v2 template | Template plasmid for cat-RNA-v2 | *Not Shown* | PcymR - cat-RNA-v2-tVoigtS4 - KanR.II - pBBR1 |
| | pMJD093 - pMJD104 | 10 Rational cat-RNA v2 | Set of 10 rationally designed orthogonal cat-RNA-v2 barcodes | 2D, 4, 5 | PcymR - cat-RNA-v2-tVoigtS4 - KanR.II - pBBR1 |
| *Using orthogonal cat-RNA to record 1 byte of information within 16S rRNA* | pMJD106 | Strong - pBBR1 - B5 | Construct #1 | 6,7 | P59 - cat-RNA-v2-tVoigtS4 - KanR.II - pBBR1 |
| | pMJD107 | Medium1 - pBBR1 - B3 | Construct #2 | 6,7 | P34 - cat-RNA-v2-tVoigtS4 - KanR.II - pBBR1 |
| | pMJD108 | Medium2 - pBBR1 - B1 | Construct #3 | 6,7 | P8 - cat-RNA-v2-tVoigtS4 - KanR.II - pBBR1 |
| | pMJD109 | Weak - pBBR1 - B4 | Construct #4 | 6,7 | P51 - cat-RNA-v2-tVoigtS4 - KanR.II - pBBR1 |
| | pMJD110 | Strong - pColE1 - B10 | Construct #5 | 6,7 | P59 - cat-RNA-v2-TtVoigtS4- KanR.II - pColE1 |
| | pMJD111 | Medium1 - pColE1 - B8 | Construct #6 | 6,7 | P34 - cat-RNA-v2-tVoigtS4 - KanR.II - pColE1 |
| | pMJD112 | Medium2 - pColE1 - B7 | Construct #7 | 6,7 | P8 - cat-RNA-v2-tVoigtS4 - KanR.II - pColE1 |
| | pMJD113 | Weak - pColE1 - B9 | Construct #8 | 6,7 | P51 - cat-RNA-v2-tVoigtS4 - KanR.II - pColE1 |

**Supplemental Table 2. Promoters used to transcribe cat-RNA.** The sequences of promoters used to transcribe cat-RNA-v1 and cat-RNA-v2 using plasmids that contain either a pBBR1 or ColE1 origin of replication.

| Name | Sequence |
|---|---|
| P1 | TTTTCTATCTACGTACTTGACAATGCTTGTCTATCGTGCTATAATCCCCGC GGCTCTACCTTAGTTTGTACGTT |
| P8 | TTTTCTATCTACGTACTTTGTGATGCTTGTCTATCGTGCTATAATCCCCGC GGCTCTACCTTAGTTTGTACGTT |
| P18 | TTTTCTATCTACGTACTTTGTGATGCTTGTCTATCGTGCTATAATCCCCGC GGCTCTACCTTAGTTTGTACGTT |
| P34 | TTTTCTATCTACGTACTGTGTAATGCTTGTCTATCGTGCTATAATCCCCGC GGCTCTACCTTAGTTTGTACGTT |
| P51 | TTTTCTATCTACGTACGGAATCATGCTTGTCTATCGTGCTATAATCCCCGC GGCTCTACCTTAGTTTGTACGTT |
| P59 | TTTTCTATCTACGTACTGGAGTATGCTTGTCTATCGTGCTATAATCCCCGC GGCTCTACCTTAGTTTGTACGTT |
| PcymR | AACAAACAGACAATCTGGTCTGTTTGTATTATGGAAAATTTTTCTGTATAAT AGATTCAACAAACAGACAATCTGGTCTGTTTGTATTAT |

**Supplemental Table 3. Sequence variation generated within cat-RNA barcodes.** With both types of cat-RNA, the sequence variation was introduced as shown in Figure 1.

| ribozyme | Name | Figure(s) | Sequence | Construct(s) |
|---|---|---|---|---|
| cat-RNA-v1 | V1-A | Fig. 2 | TTGGCGCCCTACTC | pSPL001,pSPL013 |
| cat-RNA-v1 | V1-B | Fig. 2 | TGGACGGACCGTAA | pSPL002,pSPL014 |
| cat-RNA-v1 | V1-C | Fig. 2 | TGGTTAGCCCAGTC | pSPL003,pSPL015 |
| cat-RNA-v1 | V1-D | Fig. 2 | TGCCCATATCGGTG | pSPL004,pSPL016 |
| cat-RNA-v1 | V1-E | Fig. 2 | TATTTAGACCGCAG | pSPL005,pSPL017 |
| cat-RNA-v1 | V1-F | Fig. 2 | TTGCCAGTGCGAGC | pSPL006,pSPL018 |
| cat-RNA-v1 | V1-G | Fig. 2 | TGCCCAGTACGCCA | pSPL007,pSPL019 |
| cat-RNA-v1 | V1-H | Fig. 2 | TACGTGCTGCGTAG | pSPL008,pSPL020 |
| cat-RNA-v1 | V1-I | Fig. 2 | TCACCAACTCGCTG | pSPL009,pSPL021 |
| cat-RNA-v1 | V1-J | Fig. 2 | TGCCTGGGTTATCC | pSPL010,pSPL022 |
| cat-RNA-v1 | V1-K | Fig. 2 | TCCGCGCCGCGCGC | pSPL011,pSPL023 |
| cat-RNA-v1 | V1-L | Fig. 2 | TCACTGCGGTATGG | pSPL012,pSPL024 |
| cat-RNA-v2 | V2-A | Fig. 2 | ATATCTCGCC | pMJD94,pMJD105 |
| cat-RNA-v2 | V2-B | Fig. 2,4,5,6,7 | CACGAAAAAA | pMJD95,pMJD106 |
| cat-RNA-v2 | V2-C | Fig. 2,4,5,6,7 | AGACCACTAC | pMJD96,pMJD107 |
| cat-RNA-v2 | V2-D | Fig. 2 | TTCATAATGT | pMJD97 |
| cat-RNA-v2 | V2-E | Fig. 2,4,5,6,7 | TCAATACTCT | pMJD98,pMJD108 |
| cat-RNA-v2 | V2-F | Fig. 2,4,5,6,7 | ATCACCGAAG | pMJD99,pMJD110 |
| cat-RNA-v2 | V2-G | Fig. 2 | GCTATAGCAA | pMJD100,pMJD111 |
| cat-RNA-v2 | V2-H | Fig. 2,4,5,6,7 | CAATTAGTAA | pMJD101 |
| cat-RNA-v2 | V2-I | Fig. 2,4,5,6,7 | AGAGAGATAA | pMJD102,pMJD112 |
| cat-RNA-v2 | V2-J | Fig. 2,4,5,6,7 | CTTAAACATA | pMJD103,pMJD113 |
| cat-RNA-v2 | V2-K | Fig. 7 | TCGCAGCTTC | pMJD105 |
| cat-RNA-v2 | V2-L | Fig. 7 | CCCCCGAGTA | pMJD105 |
| cat-RNA-v2 | V2-M | Fig. 7 | ACCACTCCAA | pMJD106 |
| cat-RNA-v2 | V2-N | Fig. 7 | AGCGCCGTAC | pMJD107 |
| cat-RNA-v2 | V2-O | Fig. 7 | GACGCGCTAG | pMJD108 |
| cat-RNA-v2 | V2-P | Fig. 7 | AATTAGGAGA | pMJD110 |
| cat-RNA-v2 | V2-Q | Fig. 7 | GACCAATAGA | pMJD111 |
| cat-RNA-v2 | V2-R | Fig. 7 | GCAGCGACCA | pMJD106 |

| | | | | |
|---|---|---|---|---|
| cat-RNA-v2 | V2-S | Fig. 7 | TTAACACCAC | pMJD113 |
| cat-RNA-v2 | V2-T | Fig. 7 | AAACACACGG | pMJD105 |
| cat-RNA-v2 | V2-U | Fig. 7 | ATCCCTATGC | pMJD106 |
| cat-RNA-v2 | V2-V | Fig. 7 | AAAACAGCCC | pMJD107 |
| cat-RNA-v2 | V2-W | Fig. 7 | TTTCTAAGGT | pMJD108 |
| cat-RNA-v2 | V2-X | Fig. 7 | ATGACGTAGC | pMJD110 |
| cat-RNA-v2 | V2-Y | Fig. 7 | AGAGCGACGG | pMJD112 |
| cat-RNA-v2 | V2-Z | Fig. 7 | TAGTATAGCC | pMJD113 |
| cat-RNA-v2 | V2-AA | Fig. 7 | AATCCGCCCG | pMJD107 |
| cat-RNA-v2 | V2-AB | Fig. 7 | AGGACTTAGA | pMJD105 |
| cat-RNA-v2 | V2-AC | Fig. 7 | TACGGTTCAT | pMJD106 |
| cat-RNA-v2 | V2-AD | Fig. 7 | TTGTTGTTCT | pMJD107 |
| cat-RNA-v2 | V2-AE | Fig. 7 | ACCCTCGTAC | pMJD108 |
| cat-RNA-v2 | V2-AF | Fig. 7 | TGAGTCGAGT | pMJD111 |
| cat-RNA-v2 | V2-AG | Fig. 7 | AAATGTTTAC | pMJD108 |
| cat-RNA-v2 | V2-AH | Fig. 7 | TTCCGAAAGT | pMJD112 |
| cat-RNA-v2 | V2-AI | Fig. 7 | TCGAGTTATT | pMJD110 |
| cat-RNA-v2 | V2-AJ | Fig. 7 | TAAGAGCGGT | pMJD111 |
| cat-RNA-v2 | V2-AK | Fig. 7 | CTTGTTACTA | pMJD113 |

**Supplemental Table 4. The 16S rRNA sequences in barcoded-rRNA amplicons.** The 16S rRNA sequences amplified span variable regions V6 through V8, allowing for species identification following barcoding.

| Microbe | Sequence |
|---|---|
| *E. coli* MG1655 16S rRNA gene | GCAACGCGAAGAACCTTACCTGGTCTTGACATCCACGGAAGTTTTC<br>AGAGATGAGAATGTGCCTTCGGGAACCGTGAGACAGGTGCTGCAT<br>GGCTGTCGTCAGCTCGTGTTGTGAAATGTTGGGTTAAGTCCCGCAA<br>CGAGCGCAACCCTTATCCTTTGTTGCCAGCGGTCCGGCCGGGAAC<br>TCAAAGGAGACTGCCAGTGATAAACTGGAGGAAGGTGGGGATGAC<br>GTCAAGTCATCATGGCCCTTACGACCAGGGCTACACACGTGCTACA<br>ATGGCGCATACAAAGAGAAGCGACCTCGCGAGAGCAAGCGGACCT<br>CATAAAGTGCGTCGTAGTCCGGATTGGAGTCTGCAACTCGACTCCA<br>TGAAGTCGGAATCGCTAGTAATCGTGGATCAGAATGCCACGGTGAA<br>TACGTTCCCGGGCCTTGTACACACCGCCCGTCA |
| *V. natriegens* 16S rRNA gene | GCAACGCGAAGAACCTTACCTACTCTTGACATCCAGAGAACTTTTCA<br>GAGATGAATTGGTGCCTTCGGGAACTCTGAGACAGGTGCTGCATGG<br>CTGTCGTCAGCTCGTGTTGTGAAATGTTGGGTTAAGTCCCGCAACG<br>AGCGCAACCCTTATCCTTGTTTGCCAGCGAGTAATGTCGGGAACTC<br>CAGGGAGACTGCCGGTGATAAACCGGAGGAAGGTGGGGACGACGT<br>CAAGTCATCATGGCCCTTACGAGTAGGGCTACACACGTGCTACAAT<br>GGCGCATACAGAGGGCGGCCAACTTGCGAAAGTGAGCGAATCCCA<br>AAAAGTGCGTCGTAGTCCGGATTGGAGTCTGCAACTCGACTCCATG<br>AAGTCGGAATCGCTAGTAATCGTGGATCAGAATGCCACGGTGAATA<br>CGTTCCCGGGCCTTGTACACACCGCCCGTCA |
| *P. putida* 16S rRNA gene | GCAACGCGAAGAACCTTACCAGGCCTTGACATGCAGAGAACTTTCC<br>AGAGATGGATTGGTGCCTTCGGGAACTCTGACACAGGTGCTGCATG<br>GCTGTCGTCAGCTCGTGTCGTGAGATGTTGGGTTAAGTCCCGTAAC<br>GAGCGCAACCCTTGTCCTTAGTTACCAGCACGTTATGGTGGGCACT<br>CTAAGGAGACTGCCGGTGACAAACCGGAGGAAGGTGGGGATGACG<br>TCAAGTCATCATGGCCCTTACGGCCTGGGCTACACACGTGCTACAA<br>TGGTCGGTACAGAGGGTTGCCAAGCCGCGAGGTGGAGCTAATCTC<br>ACAAAACCGATCGTAGTCCGGATCGCAGTCTGCAACTCGACTGCGT<br>GAAGTCGGAATCGCTAGTAATCGCGAATCAGAATGTCGCGGTGAAT<br>ACGTTCCCGGGCCTTGTACACACCGCCCGTCACACC |

**Supplemental Table 5. Primers used for RT-qPCR and NGS.** The primer-pairs used for PCRs along with an extra oligo (either probe for qPCR or reverse transcription primer for amplicon sequencing). The data corresponding to each specific primer pair is also noted for convenience. The nucleotides noted in lower case indicate partial illumina adapter sequences for NGS sequencing or restriction enzyme binding sites.

| Category | Target of the primer pair | Figure | Forward primer | | Reverse primer | | Extra oligo (Probe / Reverse Transcription primer) | | |
|---|---|---|---|---|---|---|---|---|---|
| | | | Name | Sequence | Name | Sequence | Oligo type | Oligo name | Oligo sequence |
| | 16S rRNA (native) | *4B,6B* | qRM05 | CTAGCTGGTCTGAGAGGATG | qRM16 | TGTGCAATATTCCCCACTGC | none | n/a | n/a |
| | cat-RNA-v1 Spliced 16S rRNA | *2C,S4B* | qRM17 | AGTCGGAATCGCTAGTAATCG | qRM14 | TGTAGGTCCCGTCATCTTTG | qPCR probe | qRMpr1 U64-FAM | CGGTGAATATGGTGTTCAATGCTTTTCCC |
| | total barcode cat-RNA-v1 | *S5* | qRM18 | CGACACAATCTGTCCTTTCG | qRM19 | TTGTAGAGCTCATCCATGCC | qPCR probe | qRMpr2: barcde-SUN | CAACGAAAAGCGTGACCACATGGT |
| | cat-RNA-V1 unspliced RAM | *S6* | qMJD159 | GCTGGGAACTAATTTGTATGCG | qRM14 | TGTAGGTCCCGTCATCTTTG | qPCR probe | qSPTpr2 - FAM | GGAGTACTCGATGGTGTTCAATGCTTTTCC |
| qPCR | cat-RNA-v2 Spliced 16S rRNA | *2D,4B, 6B* | qRM17 | AGTCGGAATCGCTAGTAATCG | qMJD153 | ATTTGAACCGACGATCTTCGG | qPCR probe | qRMpr1 U64-FAM | CGGTGAATATGGTGTTCAATGCTTTTCCC |
| | cat-RNA-v1 Linearize | *Cloning* | oMJD141 | ACGACACGTCTCGgTTATCCGGATCACATGAA | oMJD142 | ACGACACGTCTCCaGGGAAAAGCATTGAACA | none | n/a | n/a |
| | cat-RNA-v1 random oligos | *Cloning* | oMJD139 | gcCGTCTCaccctNNNYRNNNYRNNNgttatGAGACGGAGCTACGGCAGTCGTAGCG | oMJD138 | CGCTACGACTGCCGTAGCTC | none | n/a | n/a |
| *Barcode insertion* | cat-RNA-v2 Linearize | *Cloning* | oMJD155 | gcGGTCTCtCGTCATCTTTGCACTCCGAC | oMJD156 | gcGGTCTCaCCGAAGATCGTCGGTTCAAATC | none | n/a | n/a |
| | native16S rRNA | *S9* | oRM20 | acactctttccctacacgacgctcttccgatctGCAACGCGAAGAACCTTACC | oRM18 | gactggagttcagacgtgtgctcttccgatcTGACGGGCGGTGWGTRCA | none | n/a | n/a |
| | spliced16S rRNA cat-RNA-v1 | *S4B,2F* | oRM20 | acactctttccctacacgacgctcttccgatctGCAACGCGAAGAACCTTACC | oRM21 | gactggagttcagacgtgtgctcttccgatctAAGTCATGCCGTTTCATGTGATC | none | n/a | n/a |
| | cat-RNA-v1 | *Not shown* | oMJD99 | acactctttccctacacgacgctcttccgatctGTTCACAGACTAAATGTCGGT | oRM18 | gactggagttcagacgtgtgctcttccgatcTGACGGGCGGTGWGTRCA | none | n/a | n/a |
| | spliced16S rRNA cat-RNA-v2 | *2G,4C-E,5,6C* | oRM20 | acactctttccctacacgacgctcttccgatctGCAACGCGAAGAACCTTACC | oMJD154 | gactggagttcagacgtgtgctcttccgatctATTTGAACCGACGATCTTCGG | none | n/a | n/a |
| *NGS* | cat-RNA-v2 | *6C* | oMJD99 | acactctttccctacacgacgctcttccgatctGTTCACAGACTAAATGTCGGT | oMJD154 | gactggagttcagacgtgtgctcttccgatctATTTGAACCGACGATCTTCGG | none | n/a | n/a |

**Supplemental Table 6. Viennafold predictions.** For each barcode (B#), statistical likelihood of properly folding, the RNA sequence of the barcode, screening for restriction enzyme and RNAse E binding sites, the DNA complement sequence, predicted Gibbs free energy of folding for the barcode sequence incorporated into the tRNA structure + the 20bp of the 16S rRNA upstream of the splice site, and the deviation from the WT structure (pLNK007).

| B# | entropy | RNA seq | Motif GUAUUU | Motif RNWUU | Motif GGTCTC | Motif GAGACC | DNA seq | MFE | Bp deviation |
|---|---|---|---|---|---|---|---|---|---|
| 1 | 0.426432 | AUAUCUCGCC | -1 | -1 | -1 | -1 | ATATCTCGCC | -43.5 | 0 |
| 2 | 0.256694 | CACGAAAAAA | -1 | -1 | -1 | -1 | CACGAAAAAA | -43.1 | 0 |
| 3 | 0.570156 | AGACCACUAC | -1 | -1 | -1 | -1 | AGACCACTAC | -43.5 | 0 |
| 4 | 0.109325 | UUCAUAAUGU | -1 | -1 | -1 | -1 | TTCATAATGT | -43.6 | 0 |
| 5 | 0.094724 | UCAAUACUCU | -1 | -1 | -1 | -1 | TCAATACTCT | -43.6 | 0 |
| 6 | 0.284901 | AUCACCGAAG | -1 | -1 | -1 | -1 | ATCACCGAAG | -43.3 | 0 |
| 7 | 0.247031 | GCUAUAGCAA | -1 | -1 | -1 | -1 | GCTATAGCAA | -44.5 | 0 |
| 8 | 0.417088 | CAAUUAGUAA | -1 | -1 | -1 | -1 | CAATTAGTAA | -43.1 | 0 |
| 9 | 0.078979 | AGAGAGAUAA | -1 | -1 | -1 | -1 | AGAGAGATAA | -43.1 | 0 |
| 10 | 0.185933 | CUUAAACAUA | -1 | -1 | -1 | -1 | CTTAAACATA | -43.1 | 0 |
| 11 | 1.140613 | AUGCUGAUGA | -1 | -1 | -1 | -1 | ATGCTGATGA | -43.1 | 0 |
| 12 | 1.507929 | UCUGGUAAUC | -1 | -1 | -1 | -1 | TCTGGTAATC | -43 | 0 |
| 13 | 0.804993 | CAUACUACUA | -1 | -1 | -1 | -1 | CATACTACTA | -43.1 | 0 |
| 14 | 1.260935 | CGUCAUUCAA | -1 | -1 | -1 | -1 | CGTCATTCAA | -43.1 | 0 |
| 15 | 1.380726 | AGGUGAUAGA | -1 | -1 | -1 | -1 | AGGTGATAGA | -43.1 | 0 |
| 16 | 0.771125 | CGCCCUACAA | -1 | -1 | -1 | -1 | CGCCCTACAA | -43.1 | 0 |
| 17 | 0.68003 | GAUCUGUUAG | -1 | -1 | -1 | -1 | GATCTGTTAG | -44.2 | 0 |
| 18 | 1.081691 | UGUUCAGCUU | -1 | -1 | -1 | -1 | TGTTCAGCTT | -43.6 | 0 |
| 19 | 0.633138 | CUUUUGUUAA | -1 | -1 | -1 | -1 | CTTTTGTTAA | -43.1 | 0 |
| 20 | 0.912868 | UUCCUACUAC | -1 | -1 | -1 | -1 | TTCCTACTAC | -43 | 0 |
| 21 | 1.024777 | UCGCAGCUUC | -1 | -1 | -1 | -1 | TCGCAGCTTC | -43 | 0 |
| 22 | 0.618811 | GCAGCGACCA | -1 | -1 | -1 | -1 | GCAGCGACCA | -44.5 | 0 |
| 23 | 0.756436 | AAUCCGCCCG | -1 | -1 | -1 | -1 | AATCCGCCCG | -43.3 | 0 |
| 24 | 0.099781 | AAAUGUUUAC | -1 | -1 | -1 | -1 | AAATGTTTAC | -43.5 | 0 |
| 25 | 1.192411 | AAUGGAUAAA | -1 | -1 | -1 | -1 | AATGGATAAA | -43.1 | 0 |
| 26 | 0.701418 | UCGAGUUAUU | -1 | -1 | -1 | -1 | TCGAGTTATT | -43.6 | 0 |
| 27 | 0.405385 | UAAGAGCGGU | -1 | -1 | -1 | -1 | TAAGAGCGGT | -43.6 | 0 |
| 28 | 0.162322 | AGAUGUGUGA | -1 | -1 | -1 | -1 | AGATGTGTGA | -43.1 | 0 |
| 29 | 0.778918 | CUUGUUACUA | -1 | -1 | -1 | -1 | CTTGTTACTA | -43.1 | 0 |
| 30 | 0.892207 | CCCCCGAGUA | -1 | -1 | -1 | -1 | CCCCCGAGTA | -43.1 | 0 |
| 31 | 0.405859 | ACCACUCCAA | -1 | -1 | -1 | -1 | ACCACTCCAA | -43.1 | 0 |
| 32 | 0.695005 | AGCGCCGUAC | -1 | -1 | -1 | -1 | AGCGCCGTAC | -43.5 | 0 |
| 34 | 0.856312 | GACGCGCUAG | -1 | -1 | -1 | -1 | GACGCGCTAG | -44.2 | 0 |
| 36 | 1.286117 | GCGCUGACCA | -1 | -1 | -1 | -1 | GCGCTGACCA | -44.5 | 0 |
| 37 | 0.681322 | AAUUAGGAGA | -1 | -1 | -1 | -1 | AATTAGGAGA | -43.1 | 0 |
| 38 | 0.543592 | GACCAAUAGA | -1 | -1 | -1 | -1 | GACCAATAGA | -44.5 | 0 |
| 39 | 0.611185 | CAUUCCCCCA | -1 | -1 | -1 | -1 | CATTCCCCCA | -43.1 | 0 |

| | | | | | | | | | |
|---|---|---|---|---|---|---|---|---|---|
| 40 | 1.083669 | UUAACACCAC | -1 | -1 | -1 | -1 | TTAACACCAC | -43 | 0 |
| 41 | 0.468193 | AAACACACGG | -1 | -1 | -1 | -1 | AAACACACGG | -43.3 | 0 |
| 42 | 0.320423 | AUCCCUAUGC | -1 | -1 | -1 | -1 | ATCCCTATGC | -43.5 | 0 |
| 43 | 1.435501 | CCAACUAGCA | -1 | -1 | -1 | -1 | CCAACTAGCA | -43.1 | 0 |
| 44 | 0.175934 | AAAACAGCCC | -1 | -1 | -1 | -1 | AAAACAGCCC | -43.5 | 0 |
| 45 | 0.904966 | UUUCUAAGGU | -1 | -1 | -1 | -1 | TTTCTAAGGT | -43.6 | 0 |
| 46 | 0.115023 | AUGACGUAGC | -1 | -1 | -1 | -1 | ATGACGTAGC | -43.5 | 0 |
| 47 | 0.484284 | UCGUCUCCUU | -1 | -1 | -1 | -1 | TCGTCTCCTT | -43.6 | 0 |
| 48 | 0.978306 | AGAGCGACGG | -1 | -1 | -1 | -1 | AGAGCGACGG | -43.3 | 0 |
| 49 | 0.576202 | UAGUAUAGCC | -1 | -1 | -1 | -1 | TAGTATAGCC | -43 | 0 |
| 50 | 0.668618 | AGGACUUAGA | -1 | -1 | -1 | -1 | AGGACTTAGA | -43.1 | 0 |
| 51 | 1.130804 | UACGGUUCAU | -1 | -1 | -1 | -1 | TACGGTTCAT | -43.6 | 0 |
| 52 | 1.051434 | UUGUUGUUCU | -1 | -1 | -1 | -1 | TTGTTGTTCT | -43.6 | 0 |
| 54 | 1.557359 | UCCGAUCUAC | -1 | -1 | -1 | -1 | TCCGATCTAC | -43 | 0 |
| 55 | 0.276056 | ACCCUCGUAC | -1 | -1 | -1 | -1 | ACCCTCGTAC | -43.5 | 0 |
| 56 | 0.156513 | UAGUGCGUAU | -1 | -1 | -1 | -1 | TAGTGCGTAT | -43.6 | 0 |
| 57 | 1.676855 | UCUUUAGCUU | -1 | -1 | -1 | -1 | TCTTTAGCTT | -43.6 | 0 |
| 58 | 0.404158 | UGAGUCGAGU | -1 | -1 | -1 | -1 | TGAGTCGAGT | -43.6 | 0 |
| 60 | 0.297157 | UUCCGAAAGU | -1 | -1 | -1 | -1 | TTCCGAAAGT | -43.6 | 0 |
| 61 | 1.491669 | UGGUCAUGAU | -1 | -1 | -1 | -1 | TGGTCATGAT | -43.6 | 0 |
| 62 | 0.532307 | AUGAUCGAGC | -1 | -1 | -1 | -1 | ATGATCGAGC | -43.5 | 0 |
| 63 | 0.200846 | GCAGCAGUUA | -1 | -1 | -1 | -1 | GCAGCAGTTA | -44.5 | 0 |
| 64 | 0.236134 | GAAUAAUCGG | -1 | -1 | -1 | -1 | GAATAATCGG | -44.2 | 0 |
| 67 | 0.381715 | AAUGGAAAAC | -1 | -1 | -1 | -1 | AATGGAAAAC | -43.5 | 0 |
| 68 | 0.241787 | AAGUGUGUAC | -1 | -1 | -1 | -1 | AAGTGTGTAC | -43.5 | 0 |
| 69 | 1.261425 | GUCAUACUUG | -1 | -1 | -1 | -1 | GTCATACTTG | -44.2 | 0 |
| 70 | 0.25889 | GAAACUGACG | -1 | -1 | -1 | -1 | GAAACTGACG | -44.2 | 0 |
